# Transport and inhibition mechanism for VMAT2-mediated synaptic loading of monoamines

**DOI:** 10.1101/2023.10.27.564374

**Authors:** Yuwei Wang, Pei Zhang, Yulin Chao, Zhini Zhu, Chuanhui Yang, Zixuan Zhou, Yaohui Li, Yonghui Long, Yuehua Liu, Dianfan Li, Sheng Wang, Qianhui Qu

## Abstract

Monoamine neurotransmitters such as serotonin and dopamine are loaded by vesicular monoamine transporter 2 (VMAT2) into synaptic vesicles for storage and subsequent release in neurons. Impaired VMAT2 function underlies various neuropsychiatric diseases. VMAT2 inhibitors reserpine and tetrabenazine are used to treat hypertension, movement disorders associated with Huntington’s Disease and Tardive Dyskinesia. Despite its physiological and pharmacological significance, the structural basis underlying substrate recognition, and inhibition by varying mechanisms remains unknown. Here we present cryo-EM structures of human apo VMAT2 in addition to states bound to serotonin, tetrabenazine, and reserpine. These structures collectively capture three states, namely the lumen-facing, occluded, and cytosol-facing conformations. Notably, tetrabenazine induces a substantial rearrangement of TM2 and TM7, extending beyond the typical rocker-switch movement. These functionally dynamic snapshots, complemented by biochemical analysis, unveil the essential components responsible for ligand recognition, elucidate the proton-driven exchange cycle, and provide a framework to design improved pharmaceutics targeting VMAT2.

## Main Text

Serotonin (5-HT), dopamine (DA), and norepinephrine (NE) are major monoamine neurotransmitters that play critical roles in a variety of physiological, emotional and behavioral functions(*1–3*). Dysregulated monoaminergic neurotransmission is involved in the pathogenesis of several neurological and psychiatric disorders(*4*), such as depression, Huntington’s disease, Parkinson’s disease (PD), Alzheimer’s disease (AD), attention-deficit hyperactivity disorder (ADHD), and schizophrenia(*5*, *6*). Despite their divergent biosynthesis routes, the uptake into the presynaptic vesicles of these monoamines converge by the action of VMAT2 driven by proton gradient, which regulates the essential monoaminergic signaling in nerve system(*4*). Moreover, VMAT2 also protects the neurons from toxicants such as MPP^+^ and methamphetamine(*7*). Highlighting the importance of VMAT2 in neurophysiology, VMAT2-depleted homozygous mice display high postnatal fatality(*8*), while conditional knockout mice display stunted body size and variable deficits in behavioral changes related to movement, anxiety, motivation, and response to drugs like amphetamine(*9*). In addition, several missense VMAT2 variants have been linked to a rare infantile-onset movement disorder(*10–13*).

VMAT2 and its close paralog VMAT1 (also known as SLC18A2 and SLC18A1, respectively) belongs to the major facilitator superfamily (MFS)(*14*). Despite sharing a sequence identity of over 60%, the two VMATs differ significantly in cellular distribution, substrate recognition and pharmacological profiles(*15*). VMAT1 is primarily found in in neuroendocrine cells in the sympathetic and peripheral nervous system, whereas VMAT2 is more broadly distributed in both central and peripheral systems. Specifically, central, peripheral and enteric neurons only express VMAT2. Of note, VMAT2 co-exists with VMAT1 in adrenal glands, and its expression is induced by stress, while VMAT1 level remains constant(*16*). Both VMAT2 and VMAT1 have a similar affinity for serotonin, but VMAT2 exhibits a preference for catecholamines (e.g., DA, NE, epinephrine) with a 3-fold higher affinity, and even more so for histamine with a 30-fold higher affinity. While the competitive inhibitors reserpine (RES) and ketanserin only show a slight preference for VMAT2 over VMAT1, the non-competitive inhibitors tetrabenazine (TBZ) and its derivatives selectively target VMAT2(*15*).

Although VMAT2 holds significant physiological and pharmaceutical importance, the precise molecular mechanisms governing its recognition and transport of monoamines remains elusive. Furthermore, the distinct pharmacological effects exerted by competitive inhibitor RES and the non-competitive inhibitor TBZ warrant in-depth investigation. Here we set to address these questions by performing single-particle cryo-electron microscopy (cryo-EM) analysis on human VMAT2 complex with substrate 5-HT, inhibitors RES and TBZ.

### Structural determination assisted by ALFA-tag/nanobody

VMAT2 (55 kDa) contains a membrane domain with 12 transmembrane helices (TMs) and lacks discernable extramembrane domains (fig. S1A), posing a significant challenge for structural characterization by cryo-EM. Here we explored the feasibility of using a small and stable helical ALFA-tag (fig. S1B) in conjunction with its high-affinity nanobody, NbALFA, as a fiducial marker(*22*). We reasoned that the addition of this modestly sized duo (15 kDa) would provide an adequate fiducial reference without overwhelming the particle alignment, a challenge, based on our experience, often encountered when using larger Fabs, Pro-Macrobodies or Legobodies. To increase the success rate, we chose Pro474 and Pro489 as fusion positions based on the AlphaFold2 model(*23*), as their rigid cyclic structure was expected to restrain the ALFA-tag in a relatively fixed position. Pull-down results showed that both truncations (Residues 1-474 and 1-489) associated with NbALFA at a similar level as wildtype protein did (fig. S1C). Consequently, we proceeded with the shorter version (Residues 1-474) for cryo-EM analysis. For simplicity, this truncation is still referred to as VMAT2 hereafter.

Overexpressed wildtype VMAT2 protein was incubated with NbALFA and purified to near homogeneity using Strep affinity chromatography followed by gel filtration (fig. S1D). The particles exhibited high contrast in vitrified thin ice (fig. S1E). Consistent with the rational design, the ALFA/NbALFA signal was clearly discernable (fig. S1F). This enabled the reconstruction of cryo-EM maps of wildtype VMAT2 in various states, including the apo state (VMAT2A), the substrate bound state with serotonin/5-HT (VMAT2S), noncompetitive inhibitor TBZ-bound state (VMAT2T), and a map of VMAT2 treated with the competitive inhibitor RES (VMAT2R) (Fig. 1, and fig. S2). Of note, no discernable density could be attributed to RES molecule in VMAT2R (fig. S2). Therefore, we turned to a RES-favorable Y422C mutant which was elegantly designed by Schuldiner and colleagues(*24*). Indeed, a strong density corresponding to RES was identified in the translocation funnel of this variant (dubbed VMAT2_YC_R) (Fig. 1, and fig. S2). The well-resolved map densities permitted unambiguously model building of most regions (fig. S3).

**Fig. 1.**
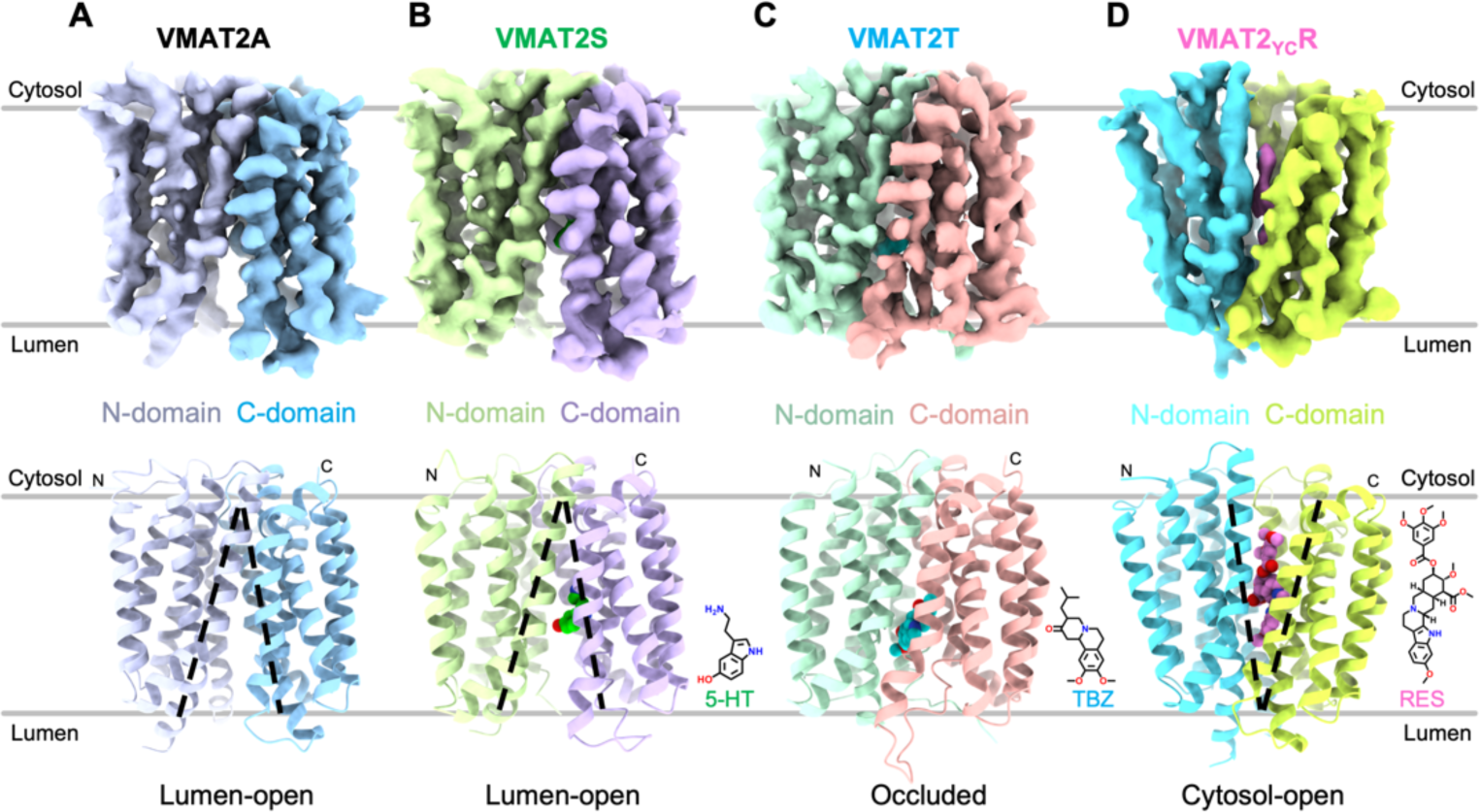
Overall structures of human VMAT2 at different ligand-bound states. (**A**) Cryo-EM density map (top) and structural model (bottom) of wildtype VMAT2 in the absence of ligand (VMAT2A), with N- and C-domain colored differently. The lumen-facing state is depicted by black dashed lines on the model. (**B**) Wildtype VMAT2 in complex with substrate serotonin (5-HT, green) captured in a lumen-facing state (VMAT2S). The 2D chemical structure of 5-HT is shown on the bottom right. (**C**) Cryo-EM density (top) and structure (bottom) of wildtype VMAT2 with non-competitive inhibitor TBZ (cyan) (VMAT2T). (**D**) Competitive inhibitor RES (pink) locks the VMAT2 Y422C mutant at a cytosol-facing state (VMAT2_YC_R).

### Overall structure and serotonin recognition

Cryo-EM map for apo VMAT2A was resolved at a nominal 3.6-Å resolution (fig. S2). VMAT2A structure adopted a canonical MFS-fold, with the N-domain composed of TMs 1-6, and the C-domain of TMs 7-12. A well-resolved intracellular linker region between TM6 and TM7 crawls along the membrane plane and connects the two pseudosymmetrically related domains. The apo structure was captured in a luminal-facing state, as judged by the large funnel opening towards the lumen space (fig. S4A), suggesting it being a lower energy resting state.

To obtain the serotonin-bound complex, we incubated the wildtype VMAT2 with 1 mM serotonin prior to vitrification. A global 3.57-Å resolution map for VMAT2S sample was obtained, with TMD local resolution higher than 3.4-Å (fig. S2). The serotonin-bound VMAT2S structure was captured in a lumen-facing state (Fig. 1B, and fig. S4B). Remarkably, despite the substrate binding, it is nearly identical to the apo structure (VMAT2A), with only a 0.6-Å Cα root mean square deviation (RMSD) (Fig. 2A). This suggests that releasing the substrate into the vesicle lumen does not require large conformational changes. Instead, several sidechains, including Glu312, Tyr341, and Phe429, shift towards the luminal space, facilitating serotonin release.

**Fig. 2.**
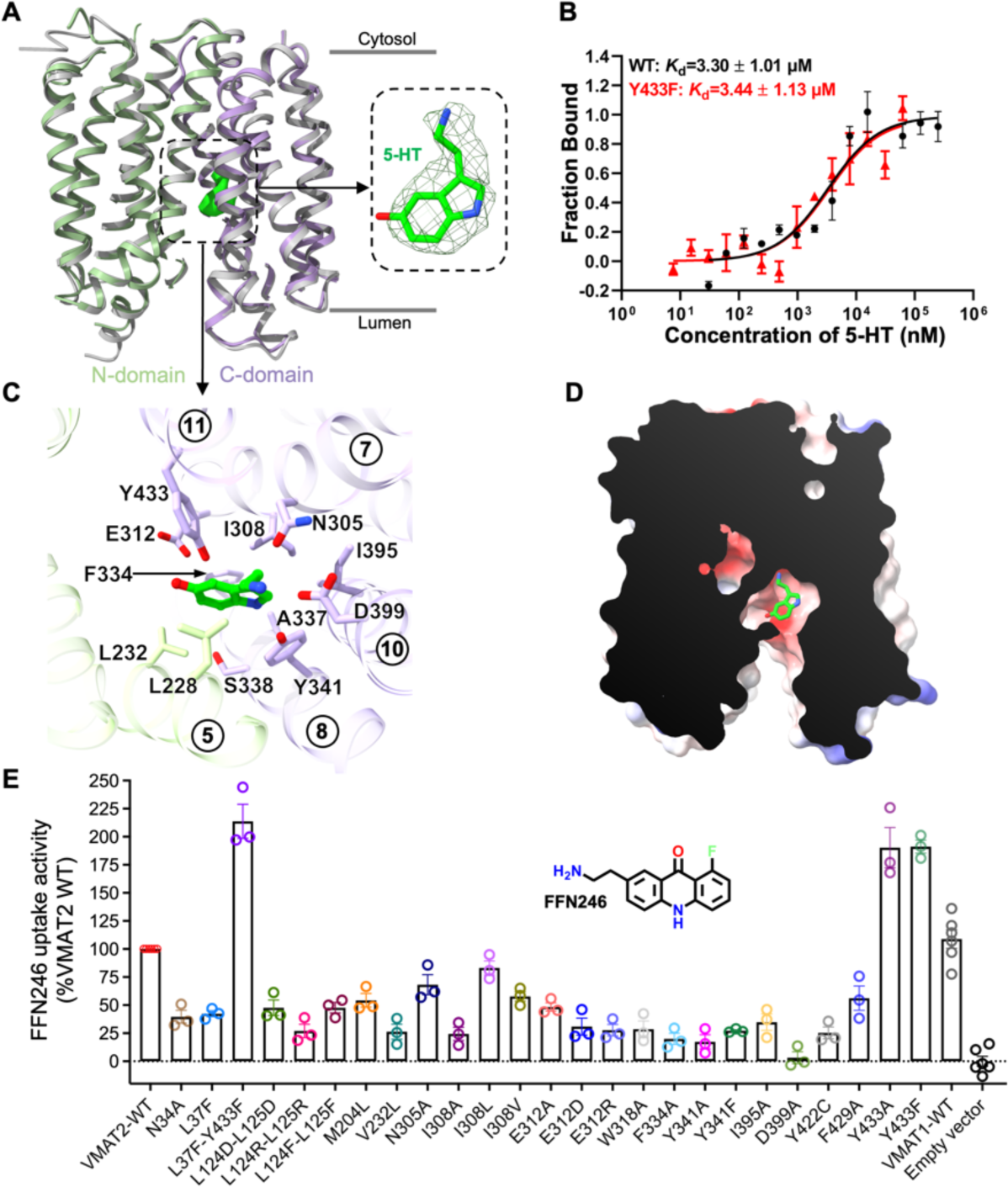
Central binding site for 5-HT in the lumen-facing state. (**A**) Structural superimposition of the lumen-facing serotonin-bound VMAT2S (N-domain in green and C-domain in purple) with apo VMAT2A (grey). The expanded view (right) shows density fitting of 5-HT (lime green). (**B**) Binding affinity for the WT and the F433F VMAT2 mutant with 5-HT measured using microscale thermophoresis assay (mean ± SEM, n=3-4 independent experiments). (**C**) Residues lining the central substrate binding cavity that accommodates 5-HT. Transmembrane helices are indicated with numbers. (**D**) Cutaway side-view of the electrostatic surface potential (negative in red, positive in blue) surface of the 5-HT binding pocket. (**E**) FFN uptake activity of VMAT2 variants. Activity values (mean ± SEM, n=3 biologically independent experiments with 3 technical replicates each) are normalized to that of the wildtype (WT).

VMAT2 features a canonical MFS translocation pathway consisting of TM1, TM4, TM5, TM7, TM8, TM10 and TM11. 5-HT was identified at the central site of the translocation pathway, as evidenced by additional density in the VMAT2S map compared to the VMAT2A map (Fig. 2A). The slightly electropositive 5-HT fits snugly within an overall electronegative cavity (Fig. 2D) surrounded by a combination of hydrophobic, polar and negatively charged residues (Fig. 2C and fig. S5). Specifically, the tryptamine group is sandwiched between Tyr341 and Phe334/Ile308, with the indole nitrogen forming an aromatic interaction with Tyr341. The primary amine of 5-HT is positioned at the negatively charged end of the pocket composed of Asp399, Tyr341 and Asn305. Additionally, the hydroxyl substitute on the 5-HT indole ring points towards the Glu312 carboxyl (Fig. 2C). In line with the observations, the binding pose was also captured in molecular dynamics simulations analysis and the molecule remained relatively stable in a duration of 500 ns (fig. S6A). In addition, mutagenic alterations to key residues lining the central binding site, such as V232L, N305A, I308A/V, E312A/D/R, Y341A/F, and D399A, drastically impaired the VMAT2-mediated uptake of FFN246, a fluorescent serotonin mimetic(*25*) (Fig. 2E). Interestingly, replacing Tyr433 with Phe or Ala only marginally weakened the binding affinity for 5-HT (Fig. 2B) but notably enhanced FFN246 transport (Fig. 2E). This suggests that a smaller side-chain at this site reduces the energetic barrier for the substate passage.

### Non-competitive inhibitor TBZ induces a unique occluded state

Originally developed as an antipsychotic drug, the non-competitive VMAT inhibitor TBZ has been used to treat movement disorders including chorea, tremor, hyperkinesia, akathisia, and tics in Europe since 1971. We incubated wildtype VMAT2 with TBZ prior to vitrification, and obtained a 3.37-Å resolution VMAT2T map.

A rod-like density in the central translocation funnel permits fitting of a TBZ molecule (Fig. 3A), which was also supported by the overall stable MD simulation trajectories (fig. S6B). It has been reported that TBZ selectively inhibits VMAT2 through the luminal opening(*26*). In our structure, VMAT2T adopts an occluded state (Fig. 3A and fig. S4C), displaying concerted movements in the luminal halves of TM1, TM2, TM7, TM8 and TM10 (Fig. 3B) compared with VMAT2A. Remarkably, beyond a rigid body rocking between N- and C-domain, a unique transformation of helix-to-loop occurred at TM2 and TM7 (fig. S4E). Specifically, the last three helical turns (Residues 313-323) of TM7 unwound at Pro313 and stretched into an unstructured loop (LuL7-8), swinging towards TM1 and TM2 of the N-domain to block the luminal exit (figs. S7A and S7B). Similarly, the luminal three helical turns of TM2 (Residues 124-132) unraveled from a hinge residue, Gly132, and ran along the newly exposed electronegative cleft after TM7’s relocation. This rearrangement latches the C-domain by buckling Leu124/Leu125 side-chains into a hydrophobic cavity occupied by TM7 apical residues Leu315 and Trp318 (figs. S7C and S7D). This reinforced occluded conformation aligns well with the proposed dead-end complex induced by TBZ(*27*, *28*).

**Fig. 3.**
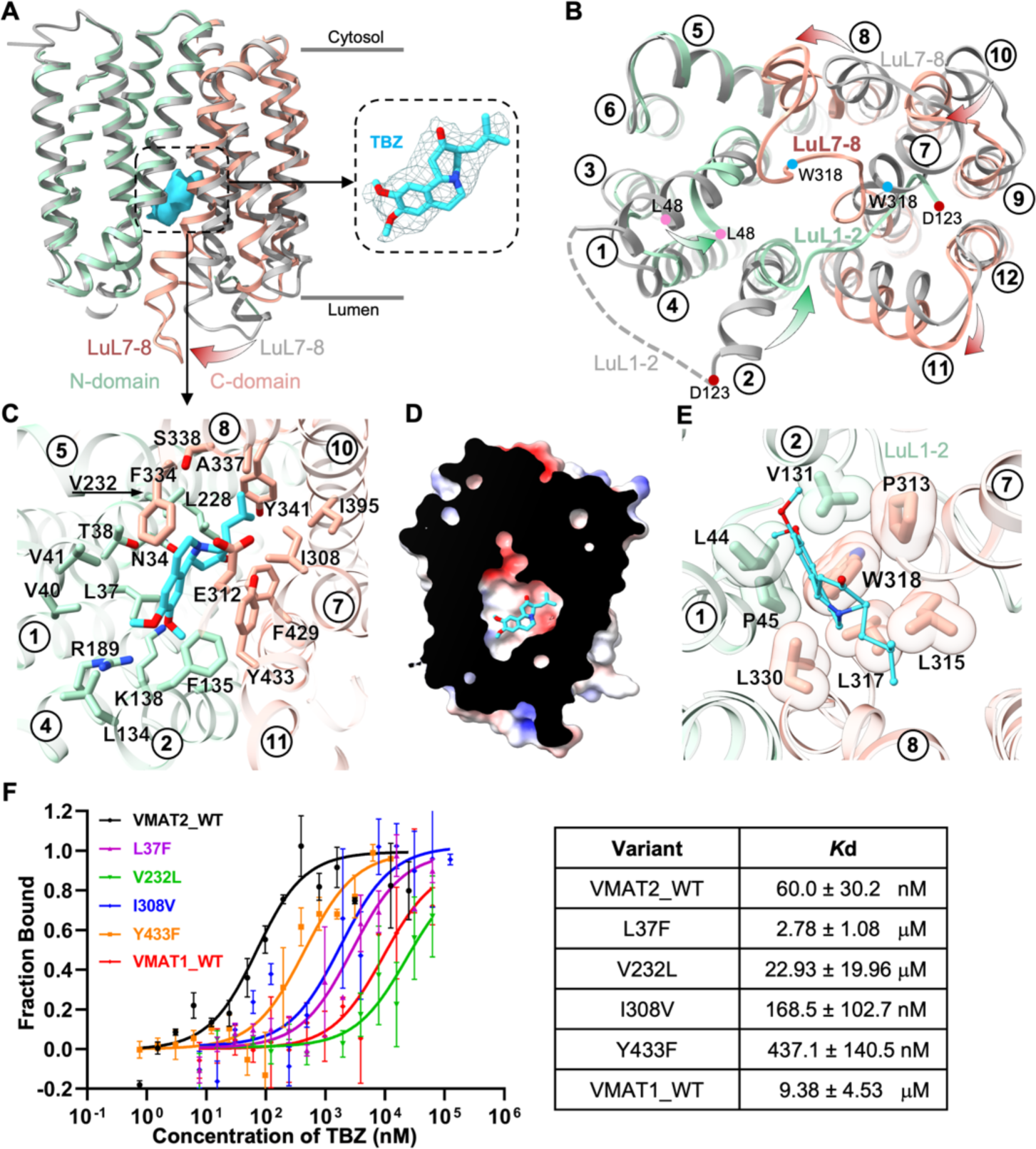
Non-competitive inhibitor TBZ locks VMAT2 in occluded state. (**A**) TBZ-bound VMAT2T structure (N-domain in light green and C-domain in salmon) is overlaid onto VMAT2A (grey). The expanded view (right) shows density fitting of TBZ (cyan). (**B**) Conformational changes induced by TBZ-binding viewed from lumen. Green and red arrows indicate movement in the N- and C-domain, respectively. Luminal loops connecting TM1/TM2 (LuL1-2) and TM7/TM8 (LuL7-8) exhibiting largest movement are labeled. (**C**) TBZ-VMAT2 interactions viewed from lumen. (**D**) Electrostatic potential surface of the hydrophobic/electronegative TBZ-binding pocket. (**E**) Residues sealing the luminal exit viewed from cytosol. (**F**) Binding affinity for VMAT2 mutants with TBZ measured using microscale thermophoresis assay. The table (right) summarizes *K*d values (mean ± SEM, *n*=3-4 independent experiments).

TBZ is situated just below the central plane of the translocation funnel (fig. S5A, middle panel). Its high binding affinity (60 nM) to VMAT2 corresponds to its occupation within an enlarged hydrophobic/electronegative pocket formed by TM1, TM2, TM4, TM5, TM7, TM8, TM10, and TM11 (Figs. 3C and 3D). TBZ is primarily nested among large aromatic residues from the C-domain and small nonpolar residues from the N-domain (Fig. S3c and fig. S5B). Substitutions including Y341A, F429A, Y433A, I308A, and I395A, had varying impacts on TBZ’s binding and inhibition activity against VMAT2 to various extents (fig. S8). This robust non-polar network positions the sole tertiary amine of TBZ close to the negatively charged Glu312 on TM7. Consistent with prior findings highlighting the critical role of Glu312 in serotonin uptake and ^3^H-TBZOH binding(*29*), the E312A mutant exhibited an 8-fold decrease in TBZ binding affinity (fig. S8A).

TBZOH (dihydrotetrabenazine) and TBZ differ only in the position of carbonyl oxygen, with TBZOH acquiring a hydroxyl group as a result of TBZ metabolism. In the TBZ-bound VMAT2 structure, the carbonyl oxygen is oriented towards Asn34 on TM1. The N34A substitution resulted in a 7-folds reduction in TBZ binding affinity (fig. S8A), suggesting a similar and important interplay between Asn34 residue and both TBZ and its metabolite TBZOH. An additional polar interaction is noted between Arg189 and the two methoxyl groups on TBZ, which may further contribute to the stable coordination network.

The intensified polar and non-polar interaction between TBZ and VMAT2 triggers the aforementioned conformational shift. Specifically, Trp318 inserts its indole group deeply towards the TBZ location from the luminal side as a result of the helix-to-loop transformation. This insertion is buttressed by several hydrophobic/non-polar residues such as Leu44, Pro45, Val131, Pro313, Leu315, Leu317, and Leu330 (Fig. 3E). While not in direct contact with TBZ in the occluded state, W318A mutant showed a substantial reduction in TBZ binding (fig. S8A). Similarly, mutagenetic perturbations on the TM2 luminal apical (Leu124 and Leu125), which is distant from the translocation funnel and TBZ binding, almost diminished TBZ association (fig. S8A). Consistently, both the W318A and L124R/L125R mutations nearly abolished the FFN uptake activity (Fig. 2E). These observations indicate that the drastic conformational change, particularly in TM2 and TM7, is critical for TBZ-induced closure of the VMAT2 luminal exit.

TBZ preferentially inhibits VMAT2, with a 10-fold lower IC_50_ value against VMAT1 (0.3 vs 3 μM)(*30*). Consistently, our *in vitro* MST assay revealed a nearly 150-fold difference in affinity between VMAT2 and VMAT1 (60 nM vs 9.38 μM, Fig. 3F). Sequence and structural comparison indicates that VMAT2 and VMAT1 differ mainly in four variable residues lining the TBZ binding pocket (fig. S9). In accordance, replacement of these VMAT2 residues with VMAT1 counterparts (L37F, V232L, I308V and Y433F) all reduced the TBZ binding to various extents (Fig. 3F). In particular, the substitution of V232L caused the utmost loss of binding affinity, consistent with previous notion that Val232 contributes the most favorable enthalpy for TBZ-VMAT2 association(*6*). In our structure, Val232 points to the TBZ isobutyl group (Fig. 3C). Extending the sidechain by a methyl group would cause clashes with the isobutyl group, explaining the weakened binding. Another important contributor is Leu37, as the L37F variant reduced TBZ binding by a substantial 45-fold, presumably also caused by steric clashes.

It is worth noting that the TBZ-bound occluded state closely resembles the predicted VMAT2 model (in the absence of any ligand) by AlphaFold2 (RMSD = 1.0 Å, fig. S4G). However, the remarkable helix-to-loop transformation and subsequent closure of luminal exit, achieved by repositioning the unwound luminal halves of TM2 and TM7, are absent in the prediction model. Our results may contribute to improving algorithms for predicting more accurate structures.

### Competitive inhibitor RES binds cytosol-facing VMAT2

Reserpine, a natural indole alkaloid, has been a first-line therapy in treating hypertension since 1955 but is currently considered as a second-line treatment due to its potential depression side effects. RES binds with high affinity to both VMAT2 and VMAT1, at the substrate-binding site on the cytoplasmic side(*30*). As mentioned earlier, we were unable to obtain the reserpine-bound complex with wildtype VMAT2 protein (fig. S2).

To capture the cytosol-facing state which was previously proposed to favor RES binding, we used the Y422C mutation (VMAT2_YC_) which was presumed to weaken the cytosolic gate(*24*). Consistently, our *in vitro* MST assay revealed that RES binds with VMAT2_YC_ mutant nearly 40-fold more strongly than to the WT (Fig. 4A). We then incubated VMAT2_YC_ with 1 mM RES and determined a of 3.7-Å structure (VMAT2_YC_R) at a cytosol-facing state (Fig. 4A, and fig. S4D). Comparing with the lumen-facing VMAT2A state, the cytosol-facing VMAT2_YC_R structure undergoes a rigid rocking-bundle movement of the N- and C-domain (fig. S4F), similar to other MFS transporters(*31*).

**Fig. 4.**
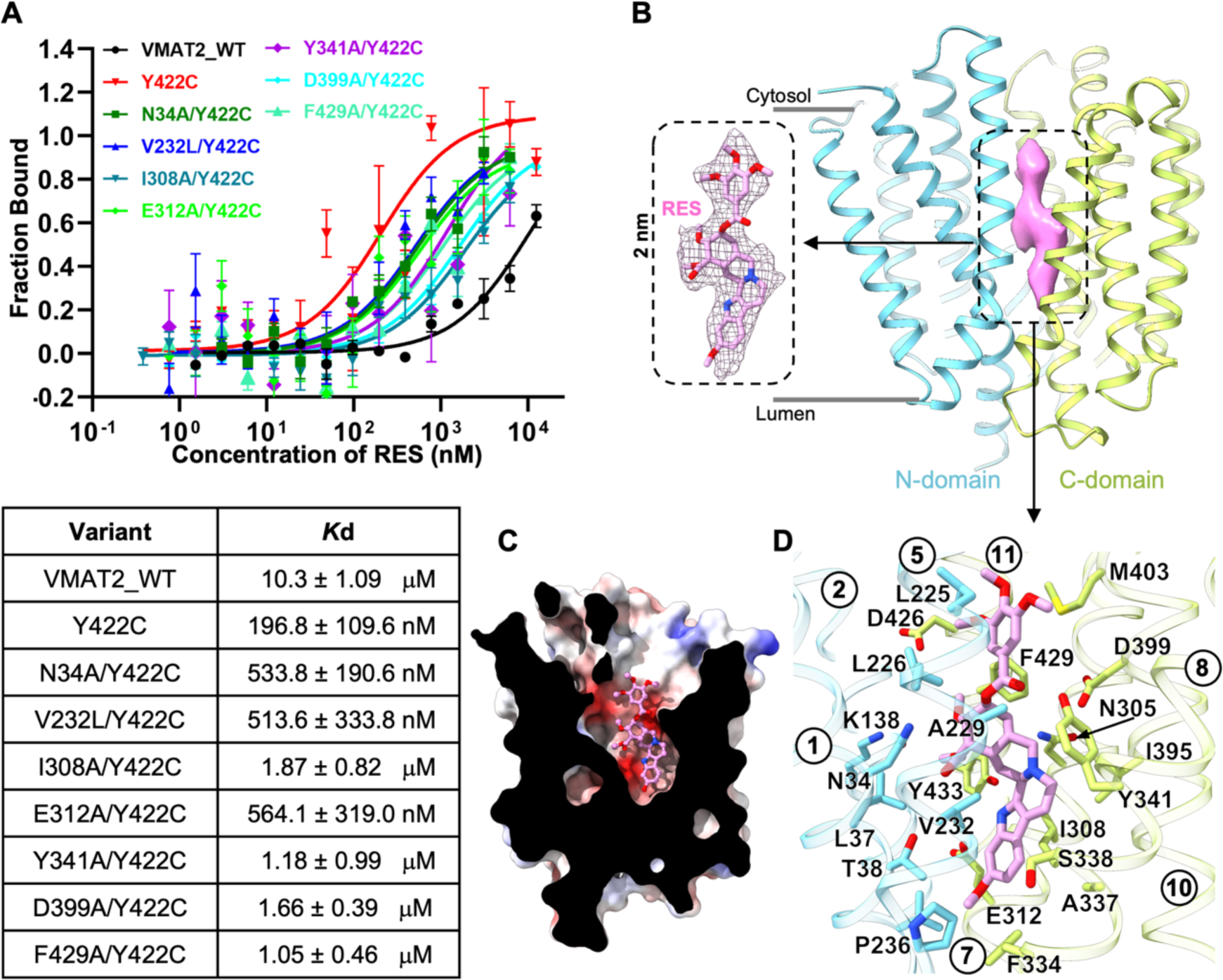
Cytosol-facing VMAT2_YC_R captured by the competitive inhibitor RES. (**A**) Binding affinity of VMAT2 variants with RES measured by MST assay, with results summarized below (mean ± SEM, n=3-4 independent experiments). (**B**) RES-bound VMAT2_YC_R structure (N-domain in cyan and C-domain in yellow green) captured at cytosol-facing state. Expanded view of the elongated density of RES (purple) is shown on left. (**C**) Cutaway side-view for the overall electronegative vestibule hosting RES molecule. (**D**) Residues in close vicinity of RES lining the translocation funnel are detailed, viewed from membrane plane.

An elongated density blob of ∼20 Å was observed in the translocation funnel of VMAT2_YC_R map. This blob matches the shape of the multi-ring reserpine (Fig. 4B). The RES trimethyoxylbenzoyl group faces the cytosol space and positions nearly perpendicular to the three adjacent rings in the middle. On the opposite side, the indole group is also perpendicular to the middle rings and wedges in the central cavity of transport passage. This indole group assumes a similar position of that of 5-HT (fig. S5A), explaining its competitive nature. The RES pose is moderately stable in MD simulation of 500 ns duration (fig. S6C). In line with our observation, substitutions on the alkaloid ring system including reserpate and reserpinediol, showed similar inhibitory efficacy on norepinephrine transport as reserpine did, while derivatives of the trimethoxylbenzoyl group inhibited neither the norepinephrine transport nor the reserpine binding(*32*).

RES binds to VMAT2 at a substantial hydrophobic/electronegative surface (Fig. 4C) lined by a combination of polar and nonpolar residues spanning nearly all TMs of the translocation funnel (Fig. 4D and fig. S5). Within the central binding site, RES orients the hydrophobic face of its methyoxyl indole group toward Val232, Pro236, Ile308, and Phe334, while directing the indole nitrogen towards Glu312 and Tyr433. In line with this binding orientation, mutations including V232L, I308A and E312A diminished RES-binding (fig. S4A). The tertiary amine on the middle tandem rings forms hydrogen bond and salt bridge with Asn305 and Asp399, respectively, as well as a cation-π interaction with Tyr341. While the N305A substitution had a minor impact on RES inhibition, D399A and Y341A/F mutations almost completely abolished FFN246 uptake (fig. S10). MST measurement further demonstrated a drastic loss of RES binding affinity in these variants (Fig. 4A), emphasizing the pivotal role of the amine-mediated interaction network in shaping the structure-activity relationship of RES. At the cytosolic vestibule, the trimethyoxylbenzoyl group brings in the cytosolic halves of the N- and C-domain by engaging with Leu225, Leu228, Met403, Ala425 and Phe429. In addition, Asp399 and Tyr341 restrict the ester linker, locking RES in position.

Notably, when the VMAT2_YC_ protein was subjected to incubation with the substrate 5-HT, the resulting reconstructed VMAT2_YC_S map adopted the same lumen-facing conformation as VMAT2S (fig. S2). These findings strongly suggest that 5-HT binding in the background of this weakened cytosolic-gate is not sufficient to transform the presumed resting lumen-facing state for substrate reception. This underscores the critical role of the cytosol-directed proton flux in driving the conformational switch of VMAT2(*28*).

### Cytosolic and luminal gates

The lumen-facing VMAT2A and the cytosol-facing VMAT2_YC_R structures provide insights into the gating mechanisms involved in the alternating access transport cycle. Akin to other MFS transporters, the cytosolic gate is composed of residues from TM4, TM5, TM10, and TM11 (Fig. 5A). In particular, the bulky Tyr422 protrudes into the middle of translocation funnel, with its aromatic face surrounded by three methionine residues (Met204, Met221 and Met403). Above this so-called Met-layer, Arg217 orchestrates an interaction network by H-bonding with the backbone oxygen of Ala208 and the sidechain hydroxyl of Tyr418, as well as a cation-ρε interaction with Tyr418. This Arg-layer effectively seals the entrance exposed to the cytosol. A unique Gly407 residue bridges the two closely stacked layers via hydrogen-bonding with the hydroxyl group of Tyr422 and the guanidine group of Arg217 (Fig. 5A). This intricate cytosolic gating network, centered around Tyr422, has been well characterized by Schuldiner and colleagues(*24*). Attenuating the gate by perturbation on Tyr422 likely lowered the energetic barrier for the conformational switch that facilitated to capture VMAT2 the cytosol-facing state.

**Fig. 5.**
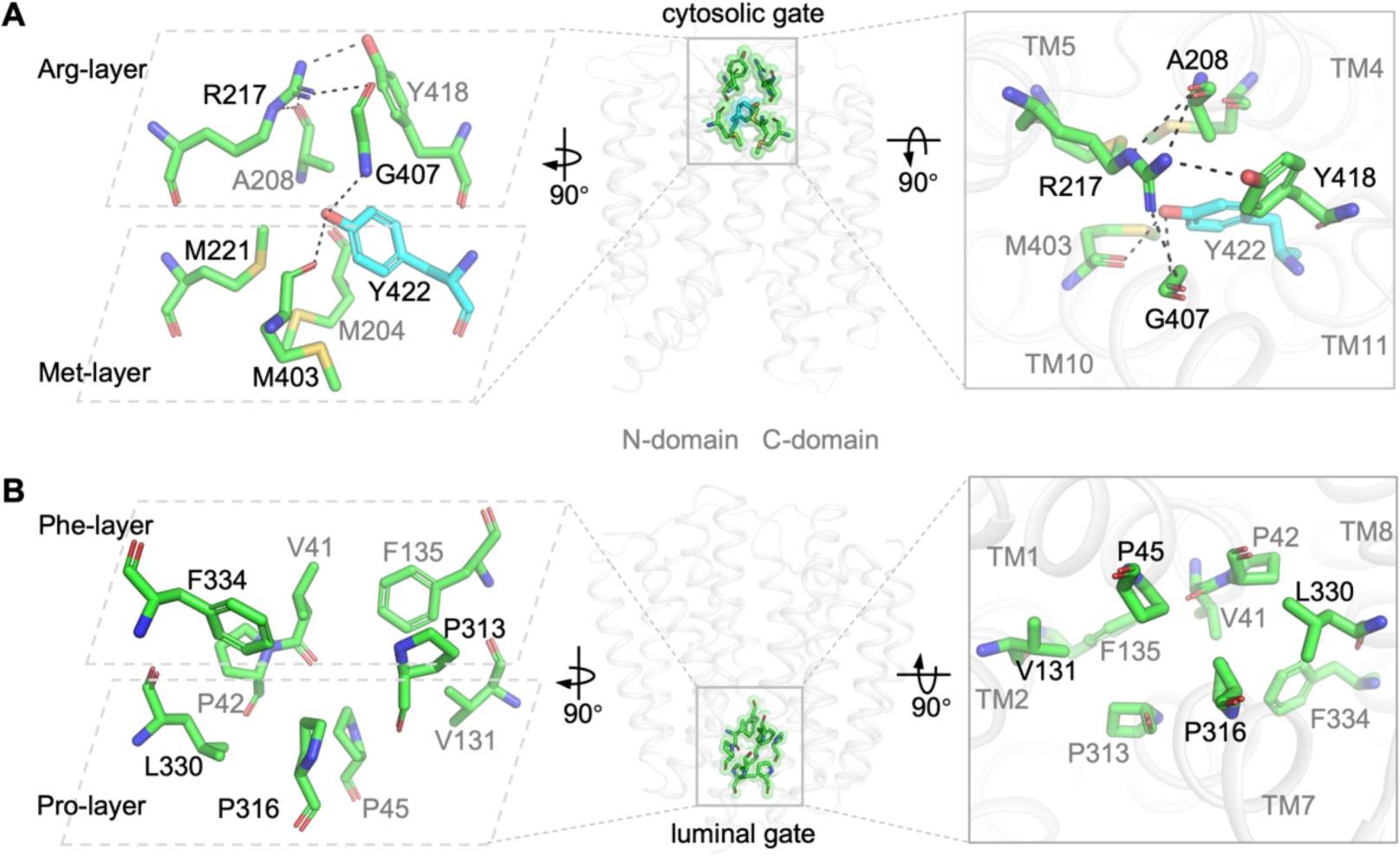
Cytosolic and luminal gates. (**A**) The cytosolic gate. Gate residues from TMs 4, 5, 10 and 11 from the VMAT2A structure (semi-transparent grey cartoon) are shown in sticks. Left, expanded view from membrane plane; right, expanded view from cytosol. The hydrophobic Met-layer and hydrophilic Arg-layer are highlighted by dashed rhomboids. (**B**) The luminal gate. Gate residues from TMs 1, 2, 7 and 8 from the cytosol-facing RES-bound VMAT2_YC_R structure are shown. Left, expanded view from membrane plane; right, expanded view from lumen. The Phe-layer and Pro-layer are highlighted.

On the luminal side, a gate is formed mainly by hydrophobic residues and prolines on TM1, TM2, TM7, and TM8 (Fig. 5B). These interactions can also be divided into two tightly packed layers. The upper Phe-layer close to the central site consists of V41, Phe135, Pro313, and Phe334, while the lower Pro-layer exposed to the lumen comprises Pro42, Pro45, Val131, Pro316 and Leu330. Substitution of Phe334 by Ala dramatically reduced transport activity (Fig. 2E). Moreover, genetic mutation of Pro316A in VMAT2 is linked to an infantile-onset form of parkinsonism(*12*). Compared with the TBZ-induced VMAT2T occluded structure, this set of luminal-gating residues in VMAT2_YC_R overlaps with the ones forming the VMAT2T luminal plug, including Pro45, Val131, and Pro313. However, the upstanding outlier in VMAT2T contributed by Trp318 blocks the exit (Fig. 3E).

### Transport mechanism

To gain more insights into the transportation cycle of VMAT2-mediated monoamine loading, we performed *in silico* molecular docking and dynamics simulation. In the cytosol-facing VMAT2 structure, 5-HT predominately adopts a single conformation, with its primary amine facing Asp399 near the cytosol side (fig. S6D). Interestingly, in the lumen-facing state, 5-HT appears in two different locations, with its primary amine pointing to either Asp399, or Glu312 near the luminal exit (fig. S6E). Experimenting the protonation state of Asp399 and Glu312 revealed a sequential binding and release of 5-HT in the translocation funnel (figs. S6F and S6G). Incorporating the general alternating access mechanism(*31*, *33*), we propose the working model for VMAT2 as follows (fig. S11).

Powered by the cytosol-directed proton gradient, VMAT2 opens its cytosolic gate (state 1) and prepares for monoamine entry (state 2). Substrate binding triggers a concerted movement of TMs (state 3) towards the synaptic lumen (state 4). In the physiological low-pH environments (∼pH 5.5), Asp399 from TM10 and Glu312 from TM7 are protonated sequentially to facilitate substrate movement within the translocation funnel and subsequent substrate release, respectively (states 5 and 6). In turn, VMAT2 is reset back (state 7) to the cytosol-open state, ready for another transport cycle (state 1). Of note, the non-competitive inhibitor tetrabenazine induces an off-cycle dead-end state through the luminal entrance.

## Conclusions

VMAT2 is responsible for packaging bioactive monoamine substances including serotonin and dopamine into presynaptic vesicles in neurons. This process primes the neurotransmitters for subsequent synaptic quantal release upon stimulation and also serve a protective mechanism for neurons against toxic substances. Our results have uncovered the structural basis for substrate recognition, and mechanism of competitive and noncompetitive inhibition by clinical drugs including reserpine and tetrabenazine. These inhibitors exploit a forgiving central binding site to achieve their potency. Along with biochemical evidence and molecular dynamic simulation analyses, our VMAT2 structures provide new insights into the mechanism underlying the vesicular packaging of monoamine neurotransmitters, offering a platform for the development of improved pharmaceutical strategies in the future.

## Materials and Methods

### Plasmids

The human full-length, wild-type VMAT2 sequence (Uniprot: Q05940) was used to create truncations (Residues 1-474 and 1-489). The full-length or truncated sequence was subcloned into a modified pcDNA3.1(-) vector, followed by an ALFA-tag, an HRV-3C site, a Twin-Strep tag, a Flag-tag, and an enhanced green fluorescence protein (eGFP). Site-directed mutagenesis was performed using homologous recombination PCR, and the sequences of all constructs were verified by DNA sequencing in Beijing Tsingke Biotech Co., Ltd.

### GST-NbALFA nanobody purification

Sequence of NbALFA nanobody was subcloned into a modified pET-21a(+) vector, with a N-terminal tag containing a GST-tag and a TEV cleavage site. For protein expression, plasmids were transformed into BL21(DE3) *E. coli* strain (Cat. TSC-E01, Beijing Tsingke Biotech Co., Ltd.). *E. coli* was cultured in terrific broth (TB) for 4 h at 37 °C before supplemented with 400 μM IPTG for 14–16 h at 18 °C. After harvest, *E. coli* cells were lysed in buffer A (50 mM HEPES, 150 mM NaCl, 5% glycerol, pH 7.4), and purified by binding to glutathione Beads (Cat. SA008100, smart-lifesciences). After extensive washing with wash buffer (50 mM HEPES, 500 mM NaCl, 5% glycerol, pH 7.4), proteins were eluted by 10 mM reduced L-Glutathione (Cat. G105426, aladdin) in buffer A. Before being incubated with VMAT2 for cryo-EM sample preparation, GST-tag was removed by using cleavage by TEV protease from nanobody. The GST-tag was spared for pull-down assay below.

### Pull down assay

To express wildtype VMAT2 and two truncations, 4 mL Expi293 cells were grown at 37 °C under a 5% CO_2_ atmosphere in Gibco® FreeStyle^TM^ 293 Expression Medium to reach the density of 2 × 10^6^ mL^-1^. 4 μg of plasmid DNA coding for recombinant wildtype VMAT2 or truncates and 8 μg PEI transfection reagent were mixed in 300 μL of medium for 15 min at RT before being added into 4 mL of cell culture. After 48 h, cells were respectively harvested by centrifugation at 1,500 g for 15 min, washed once with PBS and were resuspended with 500 μL buffer A containing 1% N-Dodecyl-β-D-maltoside (DDM, Anatrace), 0.1% cholesteryl hemisuccinate (CHS, Anatrace), 1mM Phenylmethylsulphonyl fluoride (PMSF) and 3 μg/mL protease inhibitor cocktail (Aprotinin: Pepstatin: Leupeptin(w/w) =1:1:1). The cell pellets were lysed by sonication for 2 min, agitated gently at 4 °C for 3h, and centrifuged at 12,000 rpm for 10 min. Supernatants served as input. The supernatant was incubated with 30 μL glutathione Beads bound 0.125 mg GST-nanobody at 4°C for 3h. Samples were washed with 1mL wash buffer containing 0.01% DDM and 0.001% CHS three times. Samples were mixed with 30 μL buffer A and 10 μL 4 x SDS sample-loading buffer. 20 μL of each sample and 3% v/v of the input were finally analysed by SDS-PAGE visualized via in-gel fluorescence. For negative control, the sample prepared from Expi293 cells without plasmid transfection was used.

### Recombinant protein expression and purification

For recombinant VMAT2 transient expression, 1 L of Expi293 cells were grown at 37 °C under a 5% CO_2_ atmosphere in Gibco® FreeStyle^TM^ 293 Expression Medium (ThermoFisher Scientific) to reach the density of 2 × 10^6^ mL^-1^. 1 mg of plasmid DNA coding for recombinant VMAT2 and 3 mg PEI transfection reagent were mixed in 100 mL of medium for 15 min at RT before being added into 1L of cell culture containing 2 mM sodium valproate. After 48-60h, cells were harvested by centrifugation at 1,500 g for 15 min, washed once with PBS, frozen in liquid nitrogen and stored at -80 °C. The cell pellet collected from 1 L of culture were resuspended with buffer A (50 mM HEPES, 150 mM NaCl, 5% glycerol, pH 7.4) containing 1% DDM, 0.1% CHS, 1mM Phenylmethylsulphonyl fluoride (PMSF) and 3 μg/mL protease inhibitor cocktail (Aprotinin: Pepstatin: Leupeptin(w/w) =1:1:1). The resuspended cell pellets were dounced about 30 min using a glass homogenizer, and agitated gently at 4 °C for 3h. The cell debris were removed by centrifuging at 12000rpm at 4 °C for 30min.The supernatant was incubated with 2 mL pre-equilibrated Streptactin Beads 4FF (Cat. SA053025, smart-lifesciences) and stirred gently at 4 °C for 2 h. Beads were packed into a gravity column (Cat. SLM001, smart-lifesciences) and washed with 10 column volume (CV) of buffer A containing 0.1% DDM and 0.01% CHS before being incubated with 1% lauryl maltose neopentyl glycol (LMNG, Anatrace) and 0.1% CHS in buffer A at 4 °C for 1 h. The column was washed with 30 CV of buffer A containing LMNG and CHS at a concentration ranging from (0.1% LMNG + 0.01% CHS) to (0.001% LMNG + 0.0001% CHS). Recombinant VMAT2 were eluted with 5mM D-Desthiobiotin (Cat. A1222, ChemCrus), 0.001% LMNG and 0.0001% CHS in buffer A. The eluted fraction using cleavage by HRV 3C protease (molar ratio protease: VMAT2=1:10) incubated with NbALFA nanobody (molar ratio VMAT2: nanobody=1:1.2) overnight. The pooled fractions were concentrated with a 100 kDa cut-off concentrator (Cat. UFC810096, Merck Millipore) and subjected to size-exclusion chromatography using a Superpose 6 10/300 GL column (Cat. 29-0915-96, Cytiva) equilibrated in buffer B (50 mM HEPES, 150 mM NaCl, 0.001% LMNG, 0.0001% CHS, pH 7.4). The peak fractions corresponding to purified recombinant VMAT2 were concentrated to 8 mg/mL for cryo-EM grid preparation. VMAT2_YC_ mutant was purified similarly. All proteins used to cryo-EM sample were the recombinant truncation (Residues 1-474) of VMAT2 and VMAT2_YC_. The sample was stored at -80 °C for further use.

### Microscale thermophoresis (MST)

Binding of VMAT2 and mutants to serotonin (Cat. S4244, Selleck), tetrabenazine (Cat. S1789, Selleck) and reserpine (Cat. S1601, Selleck) were measured by MST experiments. The eGFP-tagged full-length VMAT2 or VMAT2 mutants was purified by SEC in the assay buffer (50 mM HEPES, 150 mM NaCl, 0.01% DDM, 0.001% CHS, pH 7.4) as described but without cleavage. Peak fractions were pooled and diluted to 40 nM. Ligand stocks (100 mM Serotonin HCl, 100 mM Tetrabenazine and 20 mM reserpine) were diluted to the highest concentration used in assay buffer. For MST measurements, a series of 16 sequential 1:1 dilutions were prepared using assay buffer for each ligand, and each ligand dilution was mixed 1:1 with diluted protein to final protein concentration of 20 nM and final ligand concentrations in the μM to nM range. The samples were incubated for 10 min at room temperature, and then loaded into Standard Monolith Capillaries (Cat. MO-K022, NanoTemper Technologies). Measurements were carried out with a Monolith NT.115 device at 80% LED power and 40% MST power. *K*d was determined using the MO.Affinity Analysis software (version 2.3, NanoTemper Technologies, Germany) with the *K*d fit function. Capillaries displaying aggregation or adsorption were excluded. Data of at least three independently pipetted measurements were analysed and *K*d are expressed as mean ± SEM. Binding curves were plotted by GraphPad Prism Prism 9.5.1 (GraphPad Software Inc., San Diego, USA).

### FFN246 uptake assay

0.5-1 million HEK 293T (ATCC CRL-11268; mycoplasma free) cells were plated into six-well plates per well and cultured at 37℃ in 5% CO_2_. Following ∼24 hours of growth when cells reach approximately 70% confluency, the cells were transfected with WT hVMAT2-eGFP or the mutation plasmids (2μg plasmid per construct) at a ratio of 4μL polyethylenimine (PEI, 1 μg μL^−1^, Polysciences, 24765-1) per μg of DNA. The next day, cells were detached by trypsin and seeded in transparent 96-well flat bottom cell culture plate (BH, H181-96) at a density of 5*10^4^ cells/well. The cells were incubated 30min under 37 °C and then 30μL FFN246 (final concentration 1μM, BioChemPartner, BCP43741) was added for another 1h incubation. The uptake of FFN246 was terminated by three times of cold fresh PBS (with 0.2%BSA) wash. The fluorescence uptake of FFN246 was immediately recorded by flow cytometry (Thermo, Attune NxT). The gating strategy is described in fig. S12.

To characterize the inhibitory effects of TBZ or RES on VMAT2-mediated FFN246 uptake, the cells were treated similarly as above described, except that the culture medium was removed and replaced by 30μL tetrabenazine or reserpine in PBS (with 0.2% BSA, 10μM digitonin) at concentration gradients, one day after plating the cells into 96 well plate. The cells were then incubated 30min under 37℃ and 30μL FFN246 was added for another 1h incubation. The uptake was terminated by three times of cold fresh PBS (with 0.2%BSA) wash. The fluorescence uptake of FFN246 was immediately recorded by flow cytometry.

### Cryo-EM sample preparation and data collection

Negative staining was used to evaluate the protein quality. In brief, 4 µL of purified recombinant hVMAT2 or hVMAT2 mutant were applied onto glow-discharged copper grids supported by a thin layer of carbon film for 80s before negative staining by 2% (w/v) uranyl acetate solution at room temperature. The negative stained grids were examined by using FEI Talos L120C (Thermo Fisher Scientific) operated at 120 kV.

For cryo-EM grids preparation, different ligands were added at final concentration of 1 mM to the concentrated VMAT2 sample (8 mg/ml) and incubated on ice for 30min. Quantifoil Au 1.2/1.3 (300 mesh) grids were glow-discharged (12 mA for 50 s) using a PELCO easiGlo instrument (Ted Pella) before applied 2.5 μL of concentrated VMAT2 sample. These grids were blotted with filter paper for 3∼6 s (100% humidity at 4 °C) in a Vitrobot Mark IV (Thermo Fisher Scientific) and vitrified in liquid ethane at liquid nitrogen temperature. The frozen grids were transferred under cryogenic conditions and stored in liquid nitrogen for subsequent screening and cryo-EM data collection.

All datasets except VMAT2_YC_R were collected on a Titan Krios G4 cryo-electron microscope operated at 300 kV, equipped with a Falcon G4i direct electron detector with a Selectris X imaging filter (Thermo Fisher Scientific), operated with a 20 eV slit size. Movie stacks were acquired using the EPU software (Thermo Fisher Scientific) in super-resolution mode with a defocus range of −1.2 to −2.0 μm and a final calibrated pixel size of 0.932 Å. The total dose per EER (electron event representation) movie was 50 e-/Å^2^.

VMAT2_YC_R data were collected on a Titan Krios G3 transmission electron microscope (Thermo Fisher Scientific) at 300 kV, and equipped with a K3 BioQuantum direct electron detector with a Gatan GIF Quantum energy filter, operated with a 20 eV slit size. Movie stacks were acquired using serial-EM software in super-resolution mode with a defocus range of −1.2 to −2.0 μm and a final calibrated pixel size of 0.832 Å. The total dose per movie was 60 e-/Å^2^.

### Cryo-EM data processing

All datasets were processed similarly in cryoSPARC (v.3.3.2)(*35*) and RELION (v.3.1.4)(*36*). For Apo VMAT2A sample, a total of 17720 EER movies were collected. Each EER movie of 1080 frames were fractionated into 40 subgroups and beam-induced motion was corrected with a MotionCor2-like algorithm implemented in RELION. Exposure-weighted micrographs were then imported to cryoSPARC for CTF (contrast transfer function) estimation by patch CTF. Particles were blob-picked and extracted with a box size of 220 pixels, and subjected to multiple rounds of 2D classification. Several rounds of heterogeneous refinement (3D classification) were conducted using *ab initio* reference maps reconstructed with 2D averages of nice feature. The good particles were then converted for Bayesian polishing in RELION, which was subsequently imported back to cryoSPARC for one more round of *ab initio* reconstruction and several rounds of heterogeneous refinement to remove residual contaminants or poor-quality particles. Final 3.6-Å VMAT2A map from 220,731 particles by local refinement. Resolution of these maps was estimated internally in cryoSPARC by gold-standard Foushurier shell correlation using the 0.143 criterion.

For WT VMAT2 sample supplemented with substrate serotonin, tetrabenazine and reserpine, total 7527, 17914 and 5402 EER movie stacks were processed in a similar way as VMAT2A data. Final 3.57-Å VMAT2S, 3.37-Å VMAT2T and 4.75-Å VMAT2R maps were obtained from 124189, 335823 and 166931 particles.

The above procedure was also applied to 8028 K3 movies for VMAT2 Y422C mutant with reserpine, and 5148 EER movies for serotonin. Final 3.74-Å VMAT2_YC_R and 4.1-Å VMAT2_YC_S maps were determined.

### Model building and refinement

Initial VMAT2 model was retrieved from AphaFold(*23*) database which is predicted as occluded conformation (ID: AF-Q05940). The predicted model was rigid-body docked into VMAT2A cryo-EM density map in ChimeraX (v.1.6)(*37*), followed by iterative manual adjustment in COOT (v.0.9.8)(*38*) and real-space refinement in Phenix (v.1.19)(*39*). Models and geometry restraints for 5-HT, tetrabenazine, and reserpine were generated by the eLBOW tool from Phenix. The model statistics were validated by Molprobity(*40*). Sidechains that do not have well-defined density were trimmed for deposition. The final refinement statistics are provided in Table S1. Structural figures were prepared in ChimeraX or PyMOL (PyMOL Molecular Graphics SYtem, v.2.3.4, Schrödinger) (https://pymol.org/2/).

### *In silico* molecular docking

Both the lumen-facing VMAT2S and the cytosol-facing VMAT2_YC_R structures were prepared in Schrödinger (Release 2021-2) for docking. Prime (Schrödinger) was used to complete the missing side-chains and to cap the chain termini. After removal of the ligand, the protonation states and tautomers were assigned at pH 7.4 ± 0.1 using Epik(*41*) in Maestro with the OPLS3 forcefield(*42*). The docking grid was centred around the binding site, with the ligand diameter midpoint box of 20 Å on all three axes. Docking is run with the Glide standard precision (SP) scoring function(*43*).

### Molecular dynamics simulations

We performed all-atom molecular dynamics (MD) simulations in explicit solvents for VMAT2S, VMAT2T, and VMAT2_YC_R. The initial proteins include residues 15-475 for outward-open conformation, residues 18-476 for occluded conformation and residues 17-473 for inward-open conformation. The chain termini were neutralized by capping groups (acetylation and methylation) to avoid termini-charge dependent effects. PropKa was used to determine the dominant protonation state of all titratable residues at pH 7.4(*44*). The CHARMM-GUI Membrane builder module(*45*) was used to place each protein in a 1:1 POPC membrane patch with 20 Å of water above and below and 0.15 M NaCl in the solution. The final systems had ∼121 POPC lipids, ∼10,600 water molecules, and initial dimensions of 76 x 76 x 104 Å^3^. The CHARMM36m force field was adopted for lipids, proteins, sodium and chloride ions, and the TIP3P model for waters(*46*). Ligands were modeled with the CHARMM CGenFF small-molecule force field(*47*). Details of system composition are listed in Table S2.

Simulations were performed using Gromacs 2020.7(*48*). For each condition, three independent simulations were run. All systems were energy minimized and equilibrated in six steps consisting of 2.5 ns long simulations, while slowly releasing the position restrain forces acting on the Cα atoms. Initial random velocities were assigned independently to each system. Production simulations were performed for 500 ns. The Verlet neighbor list was updated every 20 steps with a cutoff of 12 Å and a buffer tolerance of 0.005 kJ/mol/ps. Non-bonded van der Waals interactions were truncated between 10 and 12 Å using a force-based switching method. Long-range electrostatic interactions under periodic boundary conditions were evaluated by using the smooth particle mesh Ewald method with a real-space cutoff of 12 Å(*49*). Bonds to hydrogen atoms were constrained with the P-LINCS algorithm with an expansion order of four and one LINCS iteration(*50*). The constant temperature was maintained at 310 K using the v-rescale (τ= 0.1 ps) thermostat(*51*) by separately coupling solvent plus salt ions, membrane, and protein. Semi-isotropic pressure coupling was applied using the Parrinello-Rahman barostat(*52*), using 1 bar and applying a coupling constant of 1 ps. Finally, a restraint-free production run was carried out for each simulation, with a time step of 2 fs.

### Data and statistical analysis

The statistical analysis was performed using GraphPad Prism 9.5.1 (GraphPad Software Inc., San Diego, USA). Binding affinity *K*d values from MST assay were calculated and graphs plotted with GraphPad Prism. The curve fitting for FFN uptake activity was determined via equation Y=Bottom + (Top-Bottom)/(1+10^((LogEC50-X))). All data are from at least three biologically independent experiments (n = 3).

## Acknowledgements

We thank the staff members of the Center of Cryo-EM of Fudan University for technical support and assistance. We thank Haochen Yang at Prof Chenqi Xu’s lab (Center of Excellence in Molecular and Cellular Science, Chinese Academy of Sciences) for guidance in flow cytometry.

## Author contributions

Q.Q. conceived the project. Y.W. and Z.Zhu purified all protein samples, and performed and analysed the biochemical assays, with assistance from Y.Li., Y.Long and H.T.. P.Z. conducted and analysed the cellular FFN246 uptake assays. C.Y. performed and analysed *in silico* docking and molecular dynamics simulations. Y.C. and Z.Zhou screened cryo-EM grids and collected cryo-EM data, with assistance of Y.W. and Z.Zhu. S.W., D.L., and Y.Liu discussed and participated in manuscript writing. Q.Q., and S.W. oversaw the project. Q.Q. wrote the manuscript with input from all authors.

## Data availability

The coordinates for apo VMAT2A, serotonin-bound VMAT2S, TBZ-bound VMAT2T and RES-bound VMAT2_YC_R have been deposited in the Protein Data Bank under accession code 8WLJ, 8WLM, 8WLK and 8WLL, respectively. The cryo-EM density maps for VMAT2A, VMAT2S, VMAT2T and VMAT2_YC_R have been deposited in the Electron Microscopy Data Bank with accession code EMD-37621, 37624, 37622 and 37623. The cryo-EM maps for VMAT2R and VMAT2_YC_S are deposited as additional maps under EMD-37623.

## References

1. J. D. Erickson, L. E. Eiden, Functional identification and molecular cloning of a human brain vesicle monoamine transporter. J. Neurochem. 61, 2314–2317 (1993).

2. D. Peter, Y. Liu, C. Sternini, R. de Giorgio, N. Brecha, R. H. Edwards, Differential expression of two vesicular monoamine transporters. J. Neurosci. Off. J. Soc. Neurosci. 15, 6179–6188 (1995).

3. E. Weihe, M. K. Schäfer, J. D. Erickson, L. E. Eiden, Localization of vesicular monoamine transporter isoforms (VMAT1 and VMAT2) to endocrine cells and neurons in rat. J. Mol. Neurosci. MN. 5, 149–164 (1994).

4. L. E. Eiden, E. Weihe, VMAT2: a dynamic regulator of brain monoaminergic neuronal function interacting with drugs of abuse. Ann. N. Y. Acad. Sci. 1216, 86–98 (2011).

5. J. K. Hsiao, W. Z. Potter, H. Agren, R. R. Owen, D. Pickar, Clinical investigation of monoamine neurotransmitter interactions. Psychopharmacology (Berl.). 112, S76–S84 (1993).

6. S. I. Støve, Å. A. Skjevik, K. Teigen, A. Martinez, Inhibition of VMAT2 by β2-adrenergic agonists, antagonists, and the atypical antipsychotic ziprasidone. Commun. Biol. 5, 1–14 (2022).

7. T. S. Guillot, G. W. Miller, Protective Actions of the Vesicular Monoamine Transporter 2 (VMAT2) in Monoaminergic Neurons. Mol. Neurobiol. 39, 149–170 (2009).

8. N. Takahashi, L. L. Miner, I. Sora, H. Ujike, R. S. Revay, V. Kostic, V. Jackson-Lewis, S. Przedborski, G. R. Uhl, VMAT2 knockout mice: Heterozygotes display reduced amphetamine-conditioned reward, enhanced amphetamine locomotion, and enhanced MPTP toxicity. Proc. Natl. Acad. Sci. 94, 9938–9943 (1997).

9. N. König, Z. Bimpisidis, S. Dumas, Å. Wallén-Mackenzie, Selective Knockout of the Vesicular Monoamine Transporter 2 (Vmat2) Gene in Calbindin2/Calretinin-Positive Neurons Results in Profound Changes in Behavior and Response to Drugs of Abuse. Front. Behav. Neurosci. 14 (2020) (available at https://www.frontiersin.org/articles/10.3389/fnbeh.2020.578443).

10. J. J. Rilstone, R. A. Alkhater, B. A. Minassian, Brain Dopamine–Serotonin Vesicular Transport Disease and Its Treatment. N. Engl. J. Med. 368, 543–550 (2013).

11. J. C. Jacobsen, C. Wilson, V. Cunningham, E. Glamuzina, D. O. Prosser, D. R. Love, T. Burgess, J. Taylor, B. Swan, R. Hill, S. P. Robertson, R. G. Snell, K. Lehnert, Brain dopamine-serotonin vesicular transport disease presenting as a severe infantile hypotonic parkinsonian disorder. J. Inherit. Metab. Dis. 39, 305–308 (2016).

12. M. Padmakumar, J. Jaeken, V. Ramaekers, L. Lagae, D. Greene, C. Thys, C. Van Geet, N. BioResource, K. Stirrups, K. Downes, E. Turro, K. Freson, A novel missense variant in SLC18A2 causes recessive brain monoamine vesicular transport disease and absent serotonin in platelets. JIMD Rep. 47, 9–16 (2019).

13. K. Saida, R. Maroofian, T. Sengoku, T. Mitani, A. T. Pagnamenta, D. Marafi, M. S. Zaki, T. J. O’Brien, E. G. Karimiani, R. Kaiyrzhanov, M. Takizawa, S. Ohori, H. Y. Leong, G. Akay, H. Galehdari, M. Zamani, R. Romy, C. J. Carroll, M. B. Toosi, F. Ashrafzadeh, S. Imannezhad, H. Malek, N. Ahangari, H. Tomoum, V. K. Gowda, V. M. Srinivasan, D. Murphy, N. Dominik, H. M. Elbendary, K. Rafat, S. Yilmaz, S. Kanmaz, M. Serin, D. Krishnakumar, A. Gardham, A. Maw, T. S. Rao, S. Alsubhi, M. Srour, D. Buhas, T. Jewett, R. E. Goldberg, H. Shamseldin, E. Frengen, D. Misceo, P. Strømme, J. R. M. Ceroni, C. A. Kim, G. Yesil, E. Sengenc, S. Guler, M. Hull, M. Parnes, D. Aktas, B. Anlar, Y. Bayram, D. Pehlivan, J. E. Posey, S. Alavi, S. A. M. Manshadi, H. Alzaidan, M. Al-Owain, L. Alabdi, F. Abdulwahab, F. Sekiguchi, K. Hamanaka, A. Fujita, Y. Uchiyama, T. Mizuguchi, S. Miyatake, N. Miyake, R. M. Elshafie, K. Salayev, U. Guliyeva, F. S. Alkuraya, J. G. Gleeson, K. G. Monaghan, K. G. Langley, H. Yang, M. Motavaf, S. Safari, M. Alipour, K. Ogata, A. E. X. Brown, J. R. Lupski, H. Houlden, N. Matsumoto, Brain monoamine vesicular transport disease caused by homozygous SLC18A2 variants: A study in 42 affected individuals. Genet. Med. 25, 90–102 (2023).

14. L. E. Eiden, M. K.-H. Schäfer, E. Weihe, B. Schütz, The vesicular amine transporter family (SLC18): amine/proton antiporters required for vesicular accumulation and regulated exocytotic secretion of monoamines and acetylcholine. Pflüg. Arch. 447, 636–640 (2004).

15. J. D. Erickson, M. K. Schafer, T. I. Bonner, L. E. Eiden, E. Weihe, Distinct pharmacological properties and distribution in neurons and endocrine cells of two isoforms of the human vesicular monoamine transporter. Proc. Natl. Acad. Sci. U. S. A. 93, 5166–5171 (1996).

16. A. Tillinger, A. Sollas, L. I. Serova, R. Kvetnansky, E. L. Sabban, Vesicular monoamine transporters (VMATs) in adrenal chromaffin cells: stress-triggered induction of VMAT2 and expression in epinephrine synthesizing cells. Cell. Mol. Neurobiol. 30, 1459–1465 (2010).

17. L. Han, Q. Qu, D. Aydin, O. Panova, M. J. Robertson, Y. Xu, R. O. Dror, G. Skiniotis, L. Feng, Structure and mechanism of the SGLT family of glucose transporters. Nature. 601, 274–279 (2022).

18. F. Li, J. Eriksen, J. Finer-Moore, R. Chang, P. Nguyen, A. Bowen, A. Myasnikov, Z. Yu, D. Bulkley, Y. Cheng, R. H. Edwards, R. M. Stroud, Ion transport and regulation in a synaptic vesicle glutamate transporter. Science. 368, 893–897 (2020).

19. Q. Zhang, X. Zhang, Y. Zhu, P. Sun, L. Zhang, J. Ma, Y. Zhang, L. Zeng, X. Nie, Y. Gao, Z. Li, S. Liu, J. Lou, A. Gao, L. Zhang, P. Gao, Recognition of cyclic dinucleotides and folates by human SLC19A1. Nature. 612, 170–176 (2022).

20. N. J. Wright, J. G. Fedor, H. Zhang, P. Jeong, Y. Suo, J. Yoo, J. Hong, W. Im, S.-Y. Lee, Methotrexate recognition by the human reduced folate carrier SLC19A1. Nature. 609, 1056– 1062 (2022).

21. B. C. McIlwain, A. L. Erwin, A. R. Davis, B. Ben Koff, L. Chang, T. Bylund, G.-Y. Chuang, P. D. Kwong, M. D. Ohi, Y.-T. Lai, R. B. Stockbridge, N-terminal Transmembrane-Helix Epitope Tag for X-ray Crystallography and Electron Microscopy of Small Membrane Proteins. J. Mol. Biol. 433, 166909 (2021).

22. H. Götzke, M. Kilisch, M. Martínez-Carranza, S. Sograte-Idrissi, A. Rajavel, T. Schlichthaerle, N. Engels, R. Jungmann, P. Stenmark, F. Opazo, S. Frey, The ALFA-tag is a highly versatile tool for nanobody-based bioscience applications. Nat. Commun. 10, 4403 (2019).

23. J. Jumper, R. Evans, A. Pritzel, T. Green, M. Figurnov, O. Ronneberger, K. Tunyasuvunakool, R. Bates, A. Žídek, A. Potapenko, A. Bridgland, C. Meyer, S. A. A. Kohl, A. J. Ballard, A. Cowie, B. Romera-Paredes, S. Nikolov, R. Jain, J. Adler, T. Back, S. Petersen, D. Reiman, E. Clancy, M. Zielinski, M. Steinegger, M. Pacholska, T. Berghammer, S. Bodenstein, D. Silver, O. Vinyals, A. W. Senior, K. Kavukcuoglu, P. Kohli, D. Hassabis, Highly accurate protein structure prediction with AlphaFold. Nature. 596, 583–589 (2021).

24. D. Yaffe, A. Vergara-Jaque, L. R. Forrest, S. Schuldiner, Emulating proton-induced conformational changes in the vesicular monoamine transporter VMAT2 by mutagenesis. Proc. Natl. Acad. Sci. 113, E7390–E7398 (2016).

25. A. Henke, Y. Kovalyova, M. Dunn, D. Dreier, N. G. Gubernator, I. Dincheva, C. Hwu, P. Šebej, M. S. Ansorge, D. Sulzer, D. Sames, Toward Serotonin Fluorescent False Neurotransmitters: Development of Fluorescent Dual Serotonin and Vesicular Monoamine Transporter Substrates for Visualizing Serotonin Neurons. ACS Chem. Neurosci. 9, 925–934 (2018).

26. D. Scherman, P. Jaudon, J. P. Henry, Characterization of the monoamine carrier of chromaffin granule membrane by binding of [2-3H]dihydrotetrabenazine. Proc. Natl. Acad. Sci. U. S. A. 80, 584–588 (1983).

27. Y. Ugolev, T. Segal, D. Yaffe, Y. Gros, S. Schuldiner, Identification of Conformationally Sensitive Residues Essential for Inhibition of Vesicular Monoamine Transport by the Noncompetitive Inhibitor Tetrabenazine. J. Biol. Chem. 288, 32160–32171 (2013).

28. D. Yaffe, L. R. Forrest, S. Schuldiner, The ins and outs of vesicular monoamine transporters. J. Gen. Physiol. 150, 671–682 (2018).

29. D. Yaffe, S. Radestock, Y. Shuster, L. R. Forrest, S. Schuldiner, Identification of molecular hinge points mediating alternating access in the vesicular monoamine transporter VMAT2. Proc. Natl. Acad. Sci. 110, E1332–E1341 (2013).

30. H. O. Lawal, D. E. Krantz, SLC18: Vesicular neurotransmitter transporters for monoamines and acetylcholine. Mol. Aspects Med. 34, 360–372 (2013).

31. N. Yan, Structural Biology of the Major Facilitator Superfamily Transporters. Annu. Rev. Biophys. 44, 257–283 (2015).

32. R. Parti, E. D. Özkan, G. J. Harnadek, D. Njus, Inhibition of Norepinephrine Transport and Reserpine Binding by Reserpine Derivatives. J. Neurochem. 48, 949–953 (1987).

33. D. Drew, R. A. North, K. Nagarathinam, M. Tanabe, Structures and General Transport Mechanisms by the Major Facilitator Superfamily (MFS). Chem. Rev. 121, 5289–5335 (2021).

34. M. P. Dalton, M. H. Cheng, I. Bahar, J. A. Coleman, BioRxiv Prepr. Serv. Biol., in press, doi:10.1101/2023.09.05.556211.

35. A. Punjani, J. L. Rubinstein, D. J. Fleet, M. A. Brubaker, cryoSPARC: algorithms for rapid unsupervised cryo-EM structure determination. Nat. Methods. 14, 290–296 (2017).

36. J. Zivanov, T. Nakane, B. O. Forsberg, D. Kimanius, W. J. Hagen, E. Lindahl, S. H. Scheres, New tools for automated high-resolution cryo-EM structure determination in RELION-3. eLife. 7 (2018), doi:10.7554/eLife.42166.

37. T. D. Goddard, C. C. Huang, E. C. Meng, E. F. Pettersen, G. S. Couch, J. H. Morris, T. E. Ferrin, UCSF ChimeraX: Meeting modern challenges in visualization and analysis. Protein Sci. Publ. Protein Soc. 27, 14–25 (2018).

38. P. Emsley, B. Lohkamp, W. G. Scott, K. Cowtan, Features and development of Coot. Acta Crystallogr. D Biol. Crystallogr. 66, 486–501 (2010).

39. P. D. Adams, P. V. Afonine, G. Bunkóczi, V. B. Chen, I. W. Davis, N. Echols, J. J. Headd, L.-W. Hung, G. J. Kapral, R. W. Grosse-Kunstleve, A. J. McCoy, N. W. Moriarty, R. Oeffner, R. J. Read, D. C. Richardson, J. S. Richardson, T. C. Terwilliger, P. H. Zwart, PHENIX: a comprehensive Python-based system for macromolecular structure solution. Acta Crystallogr. D Biol. Crystallogr. 66, 213–221 (2010).

40. V. B. Chen, W. B. Arendall, J. J. Headd, D. A. Keedy, R. M. Immormino, G. J. Kapral, L. W. Murray, J. S. Richardson, D. C. Richardson, MolProbity: all-atom structure validation for macromolecular crystallography. Acta Crystallogr. D Biol. Crystallogr. 66, 12–21 (2010).

41. J. C. Shelley, A. Cholleti, L. L. Frye, J. R. Greenwood, M. R. Timlin, M. Uchimaya, Epik: a software program for pKaprediction and protonation state generation for drug-like molecules. J. Comput. Aided Mol. Des. 21, 681–691 (2007).

42. E. Harder, W. Damm, J. Maple, C. Wu, M. Reboul, J. Y. Xiang, L. Wang, D. Lupyan, M. K. Dahlgren, J. L. Knight, J. W. Kaus, D. S. Cerutti, G. Krilov, W. L. Jorgensen, R. Abel, R. A. Friesner, OPLS3: A Force Field Providing Broad Coverage of Drug-like Small Molecules and Proteins. J. Chem. Theory Comput. 12, 281–296 (2016).

43. T. A. Halgren, R. B. Murphy, R. A. Friesner, H. S. Beard, L. L. Frye, W. T. Pollard, J. L. Banks, Glide: A New Approach for Rapid, Accurate Docking and Scoring. 2. Enrichment Factors in Database Screening. J. Med. Chem. 47, 1750–1759 (2004).

44. M. H. M. Olsson, C. R. Søndergaard, M. Rostkowski, J. H. Jensen, PROPKA3: Consistent Treatment of Internal and Surface Residues in Empirical pKa Predictions. J. Chem. Theory Comput. 7, 525–537 (2011).

45. E. L. Wu, X. Cheng, S. Jo, H. Rui, K. C. Song, E. M. Dávila-Contreras, Y. Qi, J. Lee, V. Monje-Galvan, R. M. Venable, J. B. Klauda, W. Im, CHARMM-GUI Membrane Builder toward realistic biological membrane simulations. J. Comput. Chem. 35, 1997–2004 (2014).

46. J. Huang, S. Rauscher, G. Nawrocki, T. Ran, M. Feig, B. L. de Groot, H. Grubmüller, A. D. MacKerell, CHARMM36m: an improved force field for folded and intrinsically disordered proteins. Nat. Methods. 14, 71–73 (2017).

47. W. Yu, X. He, K. Vanommeslaeghe, A. D. MacKerell, Extension of the CHARMM General Force Field to Sulfonyl-Containing Compounds and Its Utility in Biomolecular Simulations. J. Comput. Chem. 33, 2451–2468 (2012).

48. M. J. Abraham, T. Murtola, R. Schulz, S. Páll, J. C. Smith, B. Hess, E. Lindahl, GROMACS: High performance molecular simulations through multi-level parallelism from laptops to supercomputers. SoftwareX. 1, 19–25 (2015).

49. P. J. Steinbach, B. R. Brooks, New spherical-cutoff methods for long-range forces in macromolecular simulation. J. Comput. Chem. 15, 667–683 (1994).

50. B. Hess, P-LINCS: A Parallel Linear Constraint Solver for Molecular Simulation. J. Chem. Theory Comput. 4, 116–122 (2008).

51. G. Bussi, D. Donadio, M. Parrinello, Canonical sampling through velocity rescaling. J. Chem. Phys. 126, 014101 (2007).

52. M. Parrinello, A. Rahman, Polymorphic transitions in single crystals: A new molecular dynamics method. J. Appl. Phys. 52, 7182–7190 (1981).

