## Supplemental file for "Transport and inhibition mechanism for VMAT2-mediated synaptic loading of monoamines"

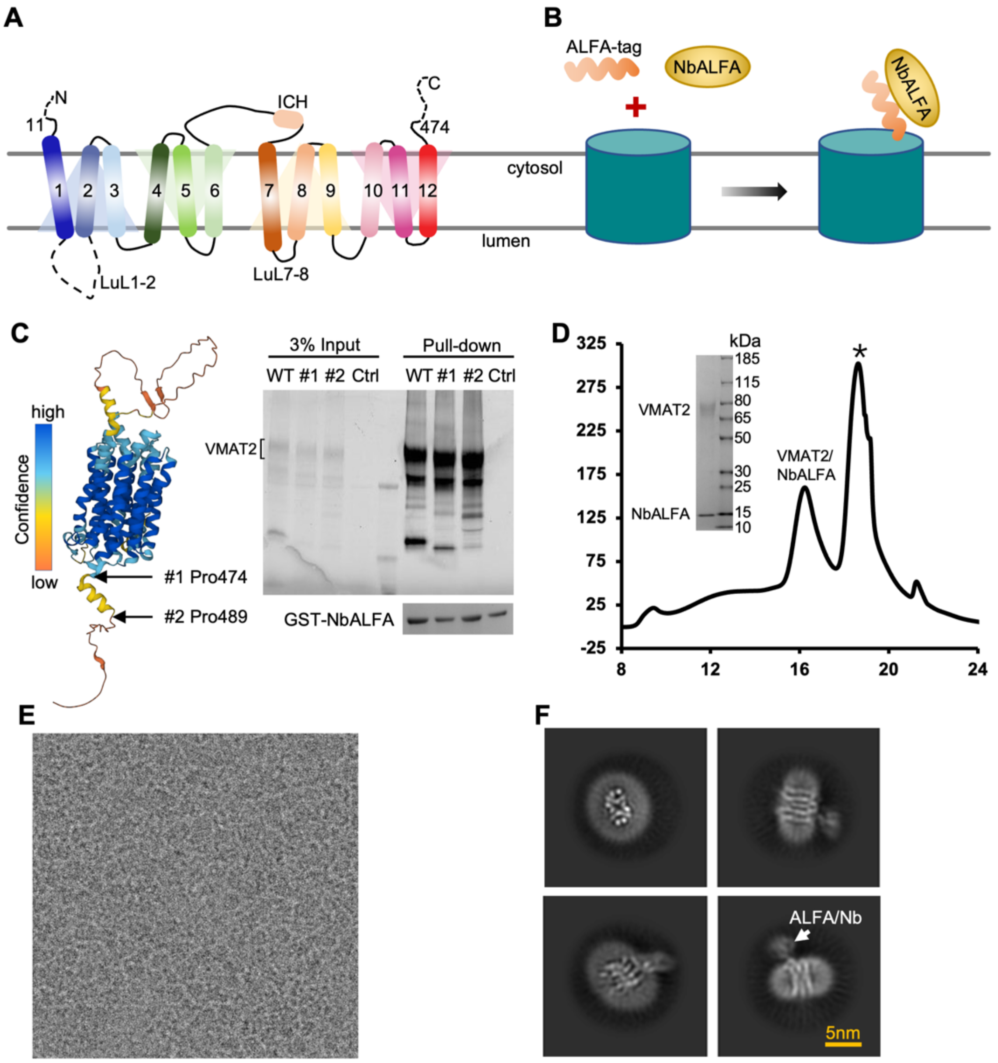


**Fig. S1. Rational design of VMAT2 fusion with helical ALFA-tag. A.** Topology diagram of full-length human VMAT2 used in the study. Regions with ill-defined densities are depicted by dashed lines. **B.** Schematic of ALFA-tag fusion and its high-affinity nanobody NbALFA as a fiducial marker for structual determination of small size membrane protein. **C.** Rational engineering based on VMAT2 AlphaFold model, with binding confirmed by pull-down assay visualized via in-gel fluorescence. **D.** Size exclusion chromatography profile and SDS-PAGE result (inset) of VMAT2-ALFA/NbALFA sample. **E.** Representative raw image of VMAT2 in thin vitrified ice. **F.** Representative 2D class averages of VMAT2 particle alignment facilitated by ALFA-tag/NbALFA marker (indicated by white arrow).

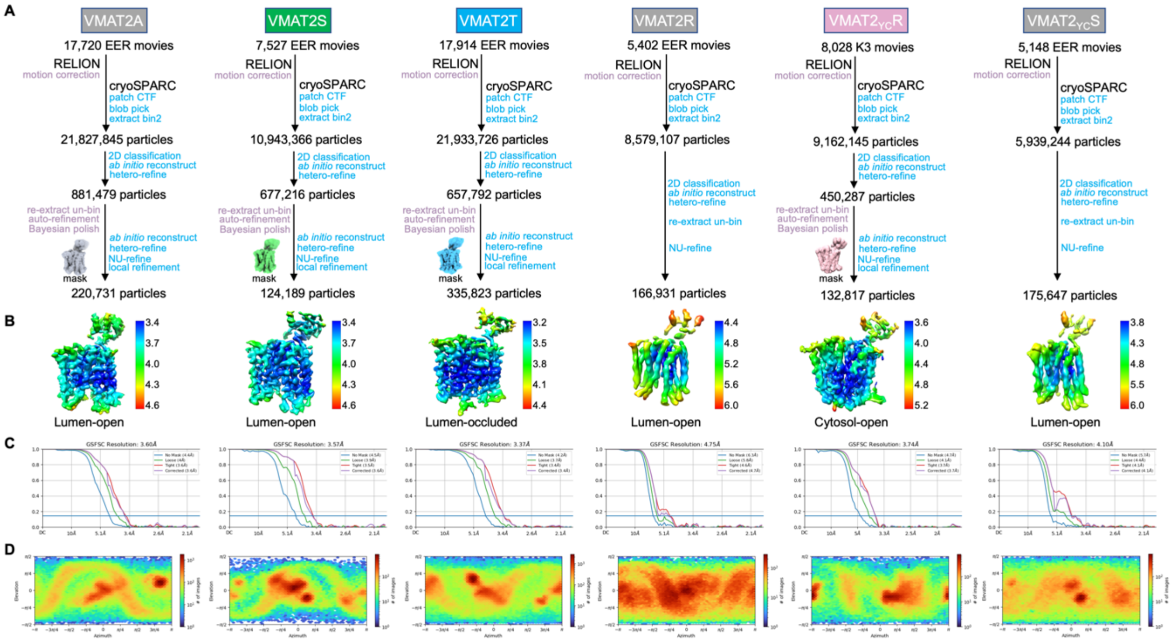


**Fig. S2. Cryo-EM analysis of VMAT2 with different ligands. A.** Processing workflow for VMAT2 wildtype protein in apo (VMAT2A), complex with substrate serotonin (VMAT2S), non-competitive inhibitor tetrabenazine (VMAT2T) and competitive inhibitor reserpine (VMAT2R), and VMAT2 Y422C mutant with reserpine ($VMAT2_{YC}R$) and serotonin ($VMAT2_{YC}S$). **B.** Final reconstructed maps colored by local resolution, with conformational states indicated below. **C.** Map resolution estimated by the gold-standard Fourier shell correlation (GSFSC) with cutoff at 0.143. **D.** Angular distribution heatmap of the final particles used in the corresponding map reconstructions.

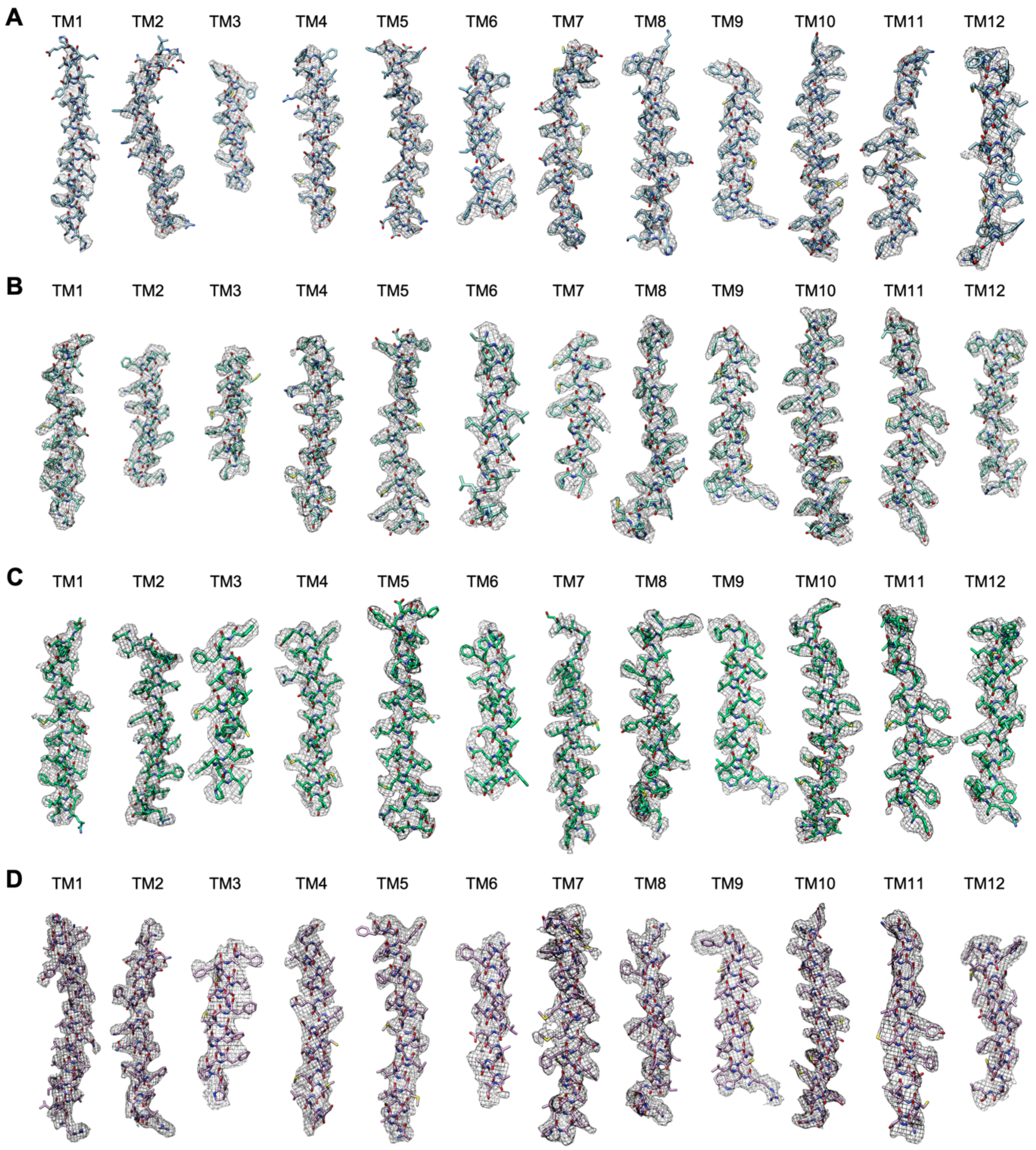


**Fig. S3. Cryo-EM density for VMAT structures.** Representative model segments fitting in the density are shown for VMAT2A (**A**), VMAT2S (**B**), VMAT2T (**C**) and $VMAT2_{YC}R$ (**D**), respectively.

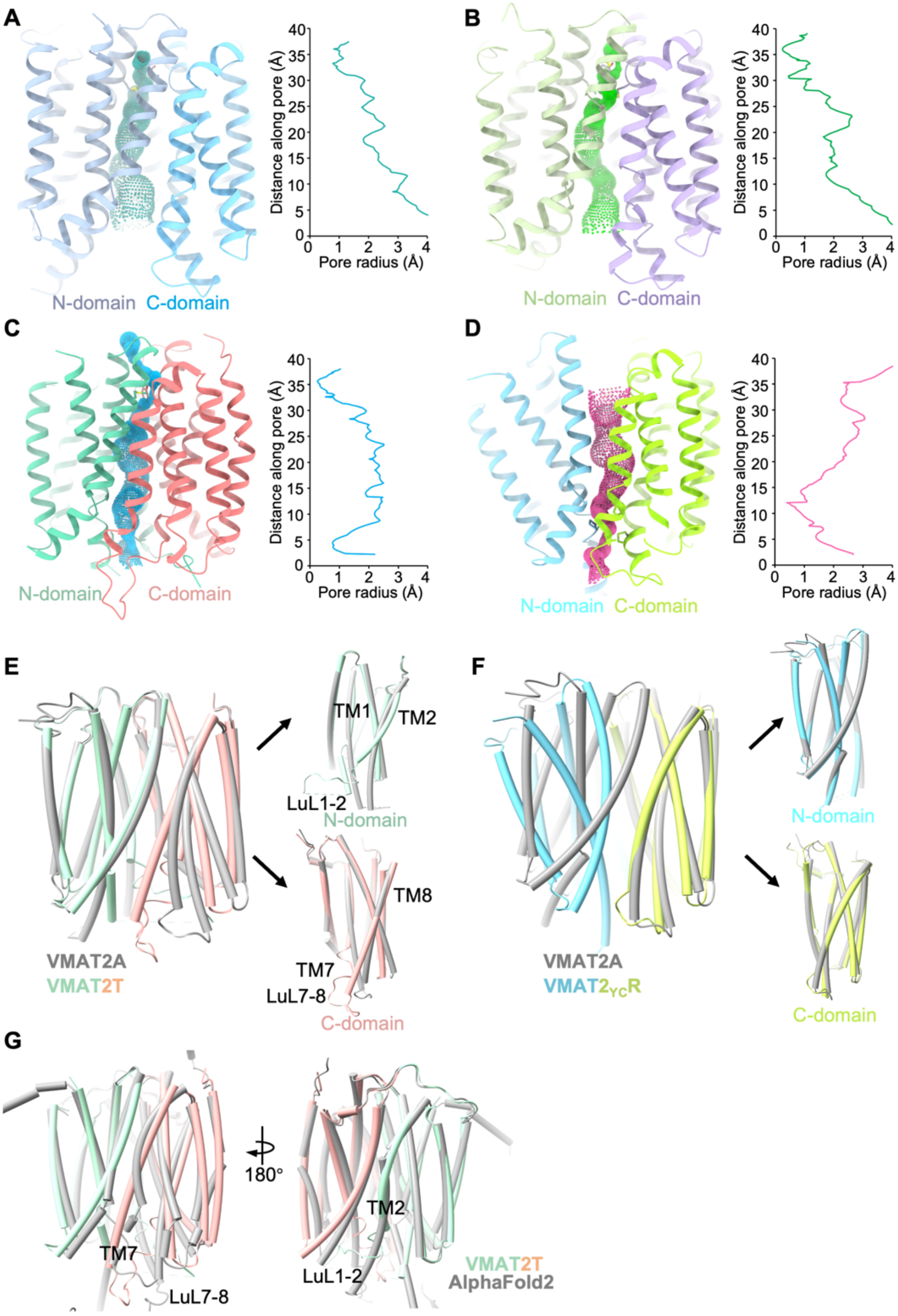


**Fig. S4. Transport funnel changes in different VMAT2 structures.** The transport pathway of VMAT2A (**A**), VMAT2S (**B**), VMAT2T (**C**) and VMAT2$_{YC}$R (**D**) is calculated by HOLE and represented in dots colored accordingly, with the corresponding radius along the transmembrane axis plotted from lumen side to cytosol. **E.** TBZ-bound VMAT2T overlaid with apo VMAT2A structure, with N- and C-domain aligned separately on the right. Significant changes of TM1-TM2 and TM7-TM8 are labelled. **F.** RES-bound VMAT2$_{YC}$R superimposed with VMAT2A, with individual N- and C-domain compared on the right. **G.** TBZ-bound VMAT2T structure is similar to the AlphaFold2 predicted model (grey), with RMSD of ~1 Å, except the unwound luminal halves of TM2 and TM7.

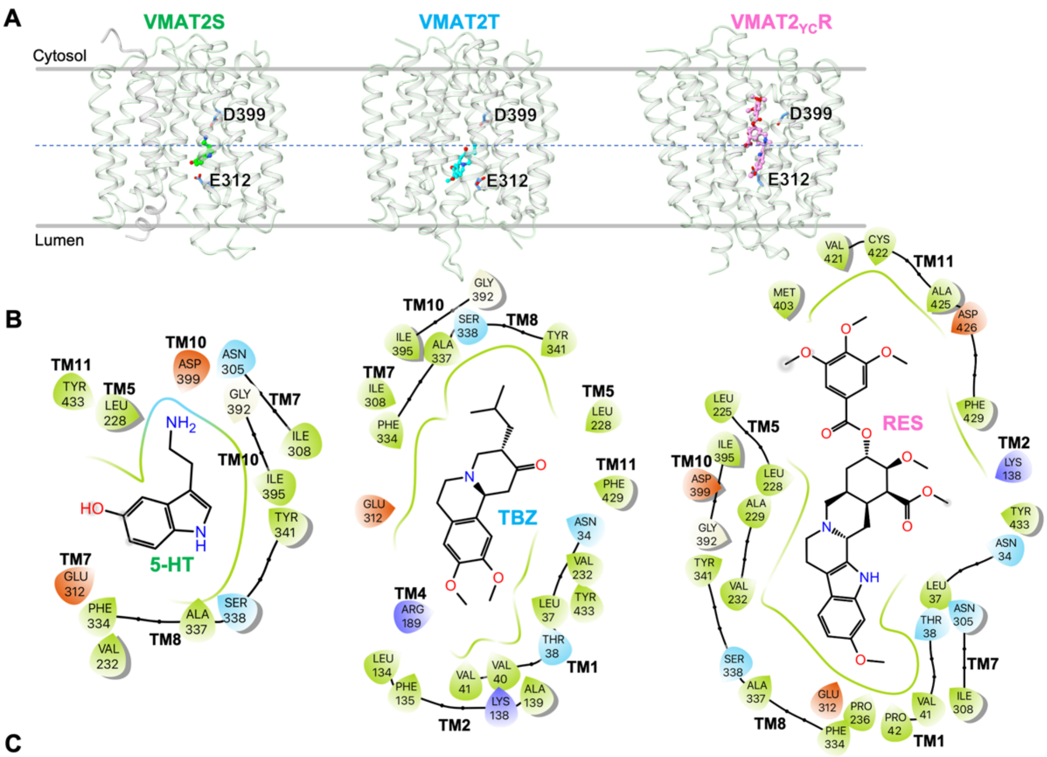


C

| N-domain | resid | 5-HT | TBZ | RES |
|---|---|---|---|---|
| TM1 | N34 | | + | + |
| | L37 | | + | + |
| | T38 | | + | + |
| | V40 | | + | |
| | V41 | | + | + |
| | P42 | | | + |
| TM2 | L134 | | + | |
| | F135 | | + | |
| | K138 | | + | + |
| | A139 | | + | |
| TM4 | R189 | | ++ | |
| TM5 | L225 | | | + |
| | L228 | + | + | + |
| | A229 | | | + |
| | V232 | + | + | + |
| | P236 | | | + |

| C-domain | resid | 5-HT | TBZ | RES |
|---|---|---|---|---|
| TM7 | N305 | + | | + |
| | I308 | + | + | + |
| | E312 | + | + | + |
| TM8 | F334 | + | + | + |
| | A337 | + | + | + |
| | S338 | + | + | + |
| | Y341 | + | + | + |
| TM10 | G392 | + | + | + |
| | I395 | + | + | + |
| | D399 | + | | + |
| | M403 | | | + |
| TM11 | V421 | | | + |
| | Y422C | | | + |
| | A425 | | | + |
| | D426 | | | + |
| | F429 | | + | + |
| | Y433 | + | + | + |

**Fig. S5. Analysis of the distinct binding patterns for 5-HT, TBZ and RES. A.** 5-HT (left) and TBZ (middle) binding sites are just below the halfway of VMAT2 translocation funnel, while RES occupies an elongated binding site across the middle point (right). **B.** 2D interaction diagram for residues interacting with 5-HT (left), TBZ (middle) and RES (right), analyzed in Maestro with a distance cutoff at 3.8 Å. Residues in green, red and blue correspond to hydrophobic, negative and positive charged features. **C.** Distribution of interaction residues in N- and C- domain of VMAT2 according to the analysis results in (**B**).

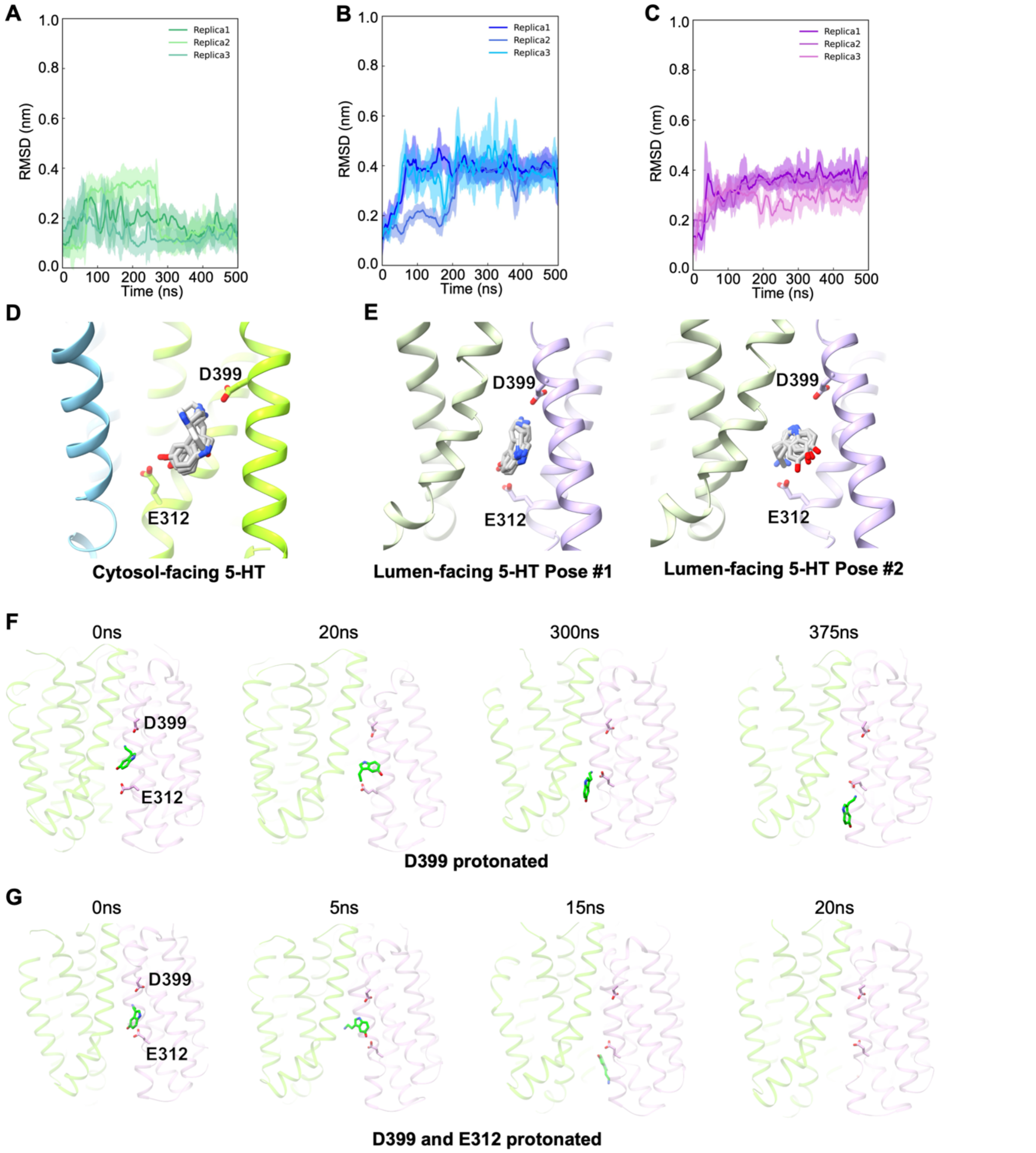


**Fig. S6. Molecular docking and dynamics simulation analysis.** Root mean square deviation (RMSD) plots of MD simulations for 5-HT (**A**), TBZ (**B**), and RES (**C**). Trajectory traces from three simulations are shown. (**D** and **E**) Five top-ranking poses from *in silico* docking of 5-HT conducted on the cytosol-facing VMAT2$_{\mathrm{YC}}$R (**D**) and the lumen-facing VMAT2S (**E**) structures. For the cytosol-facing state, nearly all 20 docking poses exhibit an invariable orientation that the primary amine points to D399 as shown in (**D**). For the lumen-facing state, 5-HT appears in two different orientations (**E**), with the primary amine facing either D399 (left) or E312 (right), of similar frequency. (**F** and **G**) MD snapshots of the 5-HT bound lumen-facing VMAT2S structure, with only D399 protonated (**F**) or both D399 and E312 protonated (**G**). Frames are selected at timepoints indicated above.

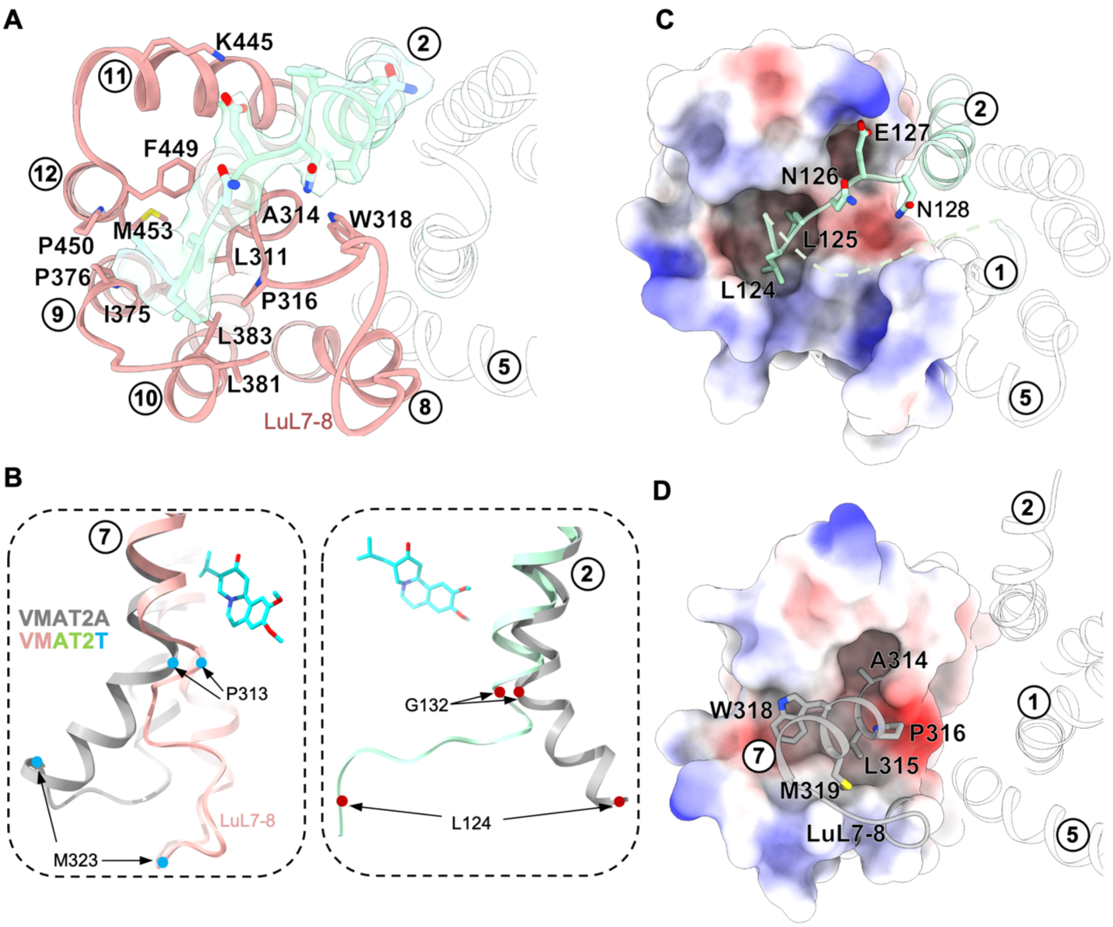


**Fig. S7. Unique conformational change induced by TBZ binding. A.** Structural alignment between apo VMAT2A (grey) and TBZ-bound VMAT2T (N-domain in green, C-domain in pink), viewed from lumen side. The luminal loop connecting TM7 and TM8 is labelled as LuL7-8, with ill-defined region depicted as dash lines. **B.** The luminal half of TM7 is unrolled and bent toward a roughly hydrophobic cavity formed by luminal ends of TMs 9, 10 and 12 in VMAT2A C-domain (shown as electrostatic surface). (**C** and **D**) The unraveled luminal half of TM2 helix is repositioned into the hydrophobic cavity occupied by TM7. Residues accommodating the uncoiling loop are shown in sticks, and the cryo-EM density in green surface representation (**C**). The reformed C-domain of VMAT2T is colored by electrostatic potential (**D**).

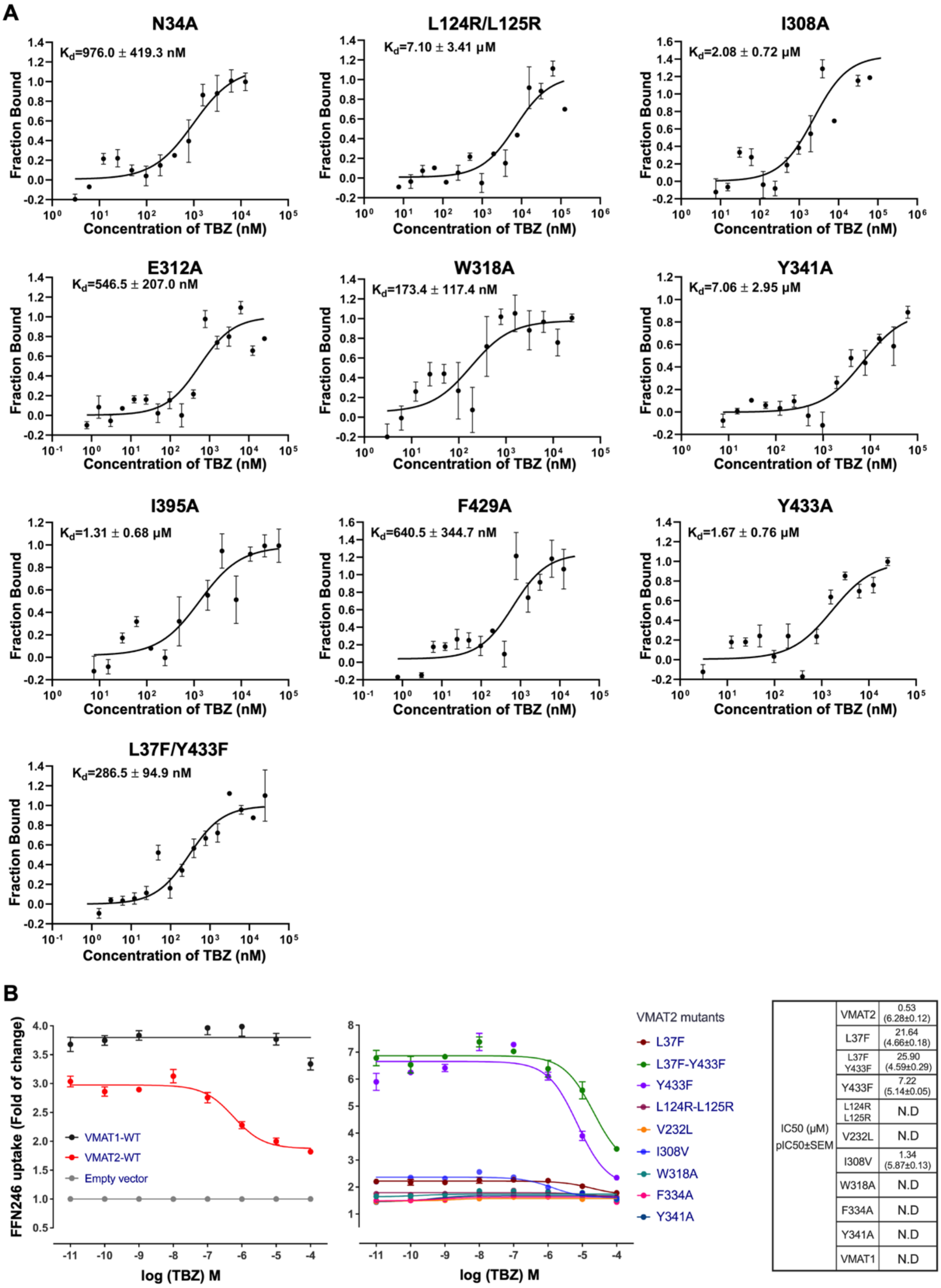


**Fig. S8. TBZ binding and FFN246 uptake of VMAT2 mutations. A.** Affinity of each VMAT2 variant is measured by microscale thermophoresis assay, and displayed with calculated *K*d value accordingly (mean ± SEM, *n*=3-4 independent experiments). **B.** FFN246 uptake activity of VMAT2 mutants affected by TBZ. Concentration-response curves for TBZ inhibition are plotted in different VMAT2 variants and WT VMAT1. $IC_{50}$ for each measurement is summarized in right table. In all panels, error bars represent SEM. N.D: the inhibitory effect of TBZ on VMAT2 mutant is not detectable.

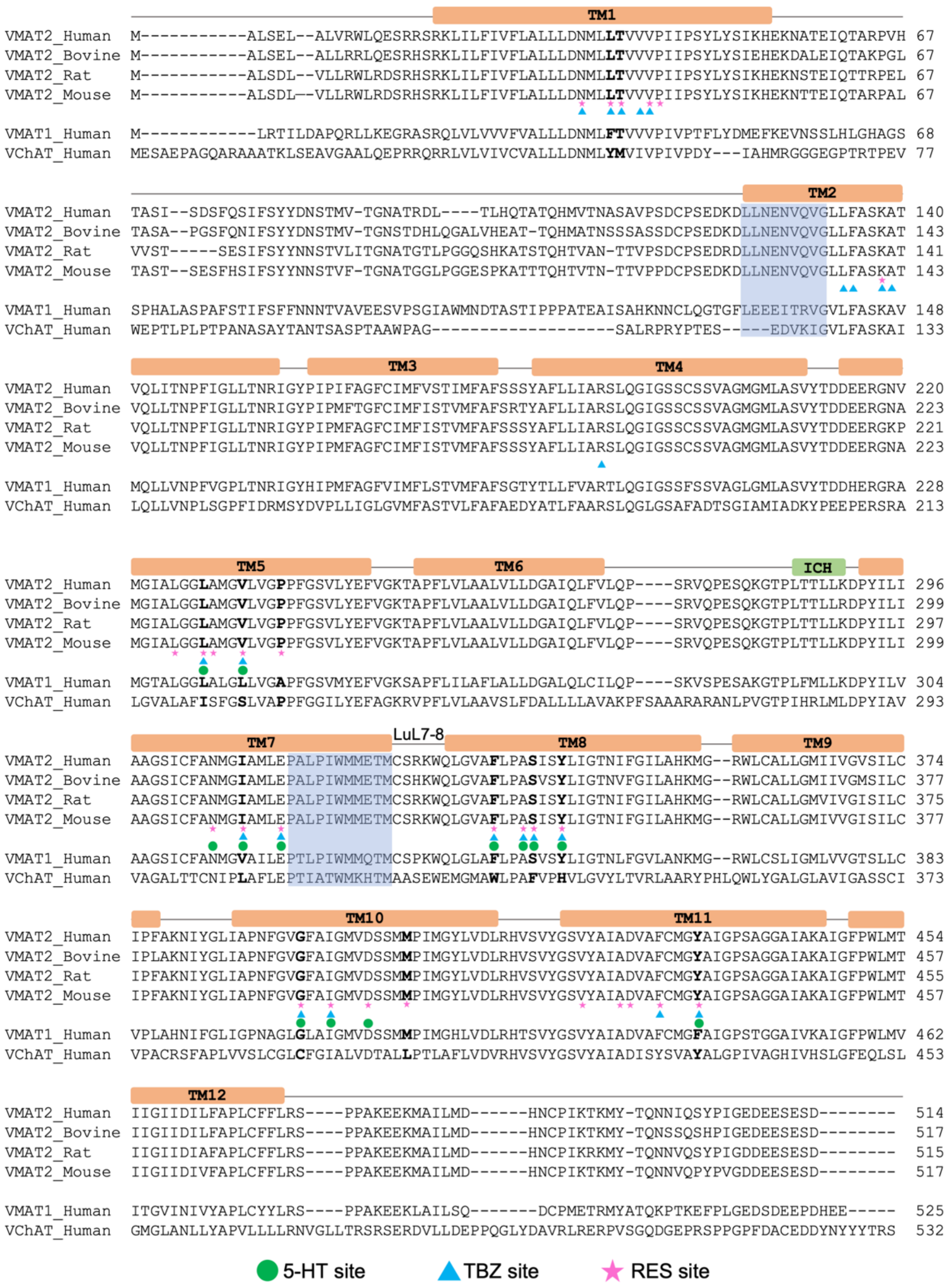


**Fig. S9. Sequence conservation of VMAT2 orthologs and human SLC18A family members.** Sequences of VMAT2/SLC18A2 from human, mouse, rat, and bovine, human VMAT1/SLC18A1, and human VChAT/SLC18A3 are compared. Secondary structure segments of human VMAT2 shown above are in accordance with Fig. 1a. Residues involved in binding substrate 5-HT, inhibitors TBZ and RES are underscored with green solid circle, cyan triangle, and pink asterisk, respectively. Ligand site residues that are not strictly conserved in VMAT1 or VChAT protein are highlighted in bold black. Helical regions of TM2 and TM7 at luminal side resolved into loops seen in TBZ-bound VMAT2T occluded state are highlighted in transparent blue shade.

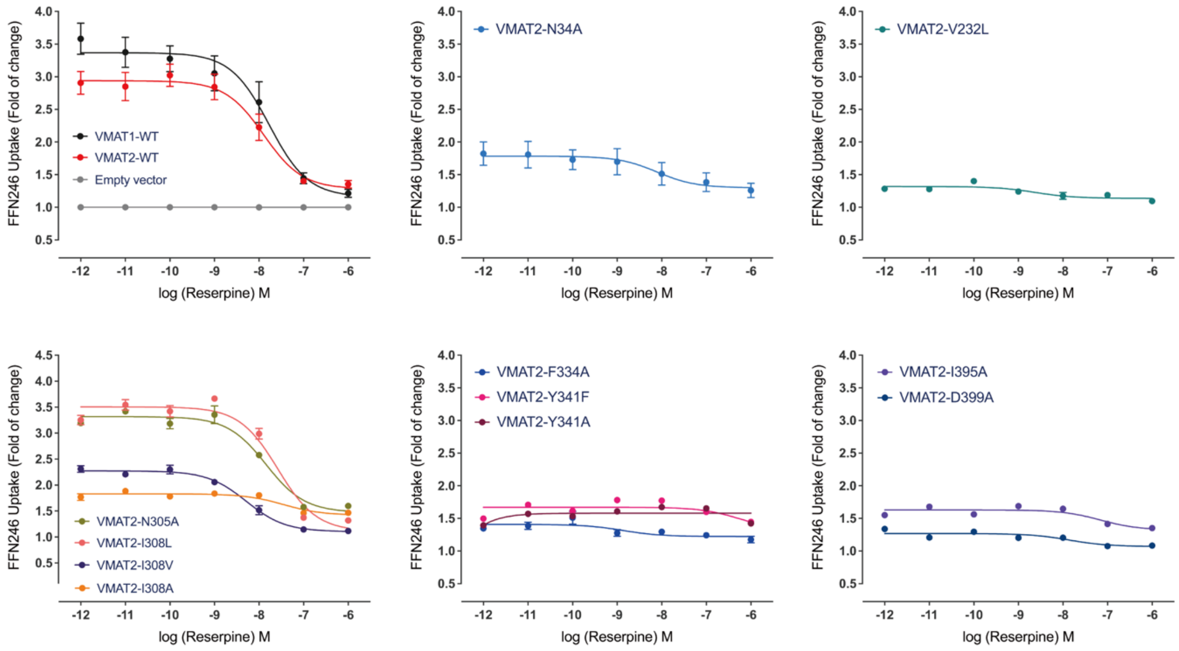


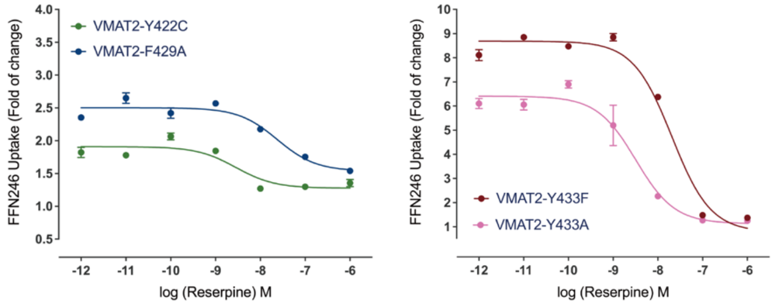


| IC50 (nM) pIC50±SEM | | | | |
|---|---|---|---|---|
| | VMAT2 | 12.27 (7.91±0.11) | Y341A | N.D |
| | N34A | 24.72 (7.61±0.23) | I395A | N.D |
| | V232L | N.D | D399A | N.D |
| | N305A | 14.38 (7.84±0.01) | Y422C | 2.82 (8.55±0.09) |
| | I308L | 26.30 (7.58±0.02) | F429A | 23.90 (7.62±0.02) |
| | I308V | 5.90 (8.23±0.18) | Y433F | 20.53 (7.69±0.02) |
| | I308A | N.D | Y433A | 2.74 (8.56±0.23) |
| | F334A | N.D | VMAT1 | 11.09 (7.95±0.24) |
| | Y341F | N.D | | |

**Fig. S10. FFN246 uptake activity of VMAT2 mutants inhibited by RES.** Concentration-response curves for reserpine inhibition are plotted in different VMAT2 variants and WT VMAT1 protein. $IC_{50}$ for each measurement is summarized in table at the right corner. In all panels, error bars represent SEM. N.D: the inhibitory effect of RES on VMAT2 mutant is not detectable.

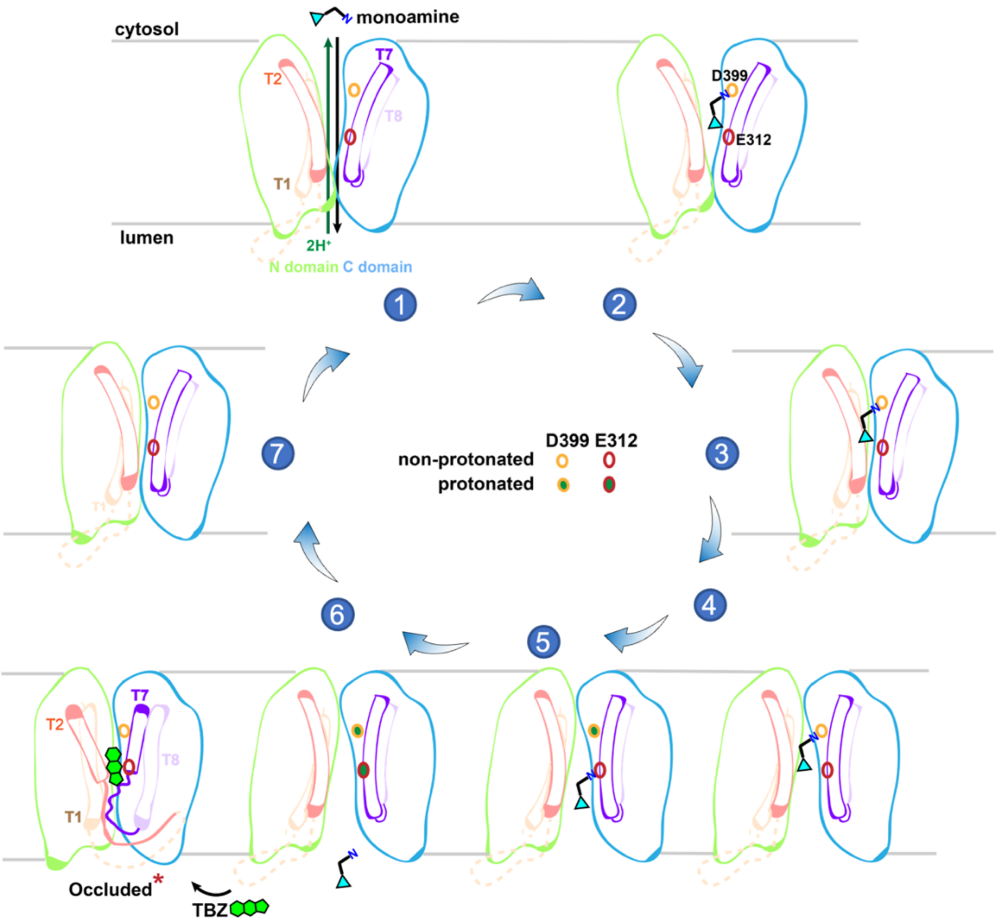


**Fig. S11. Proposed mechanism for VMAT2-mediated transport of monoamines.** Schematic representation of the alternating access transport cycle. The 7 states are derived from direct experimental structures (States 4 and 6), docking poses (States 1, 2 and 5), and AlphaFold2 prediction (States 3 and 7). For clarity, only TMs 1, 2, 7 and 8 are shown as empty tubes in color. Two negative residues D399 and E312 along the translocation pathway (empty circles, non-protonated; solid circles filled with green, protonated) facilitate substrate movement by alternating protonation states. Significant conformational shifts, particularly in TM2 and TM7, triggered by TBZ (green) entrance at the luminal side induces a dead-end occluded state.

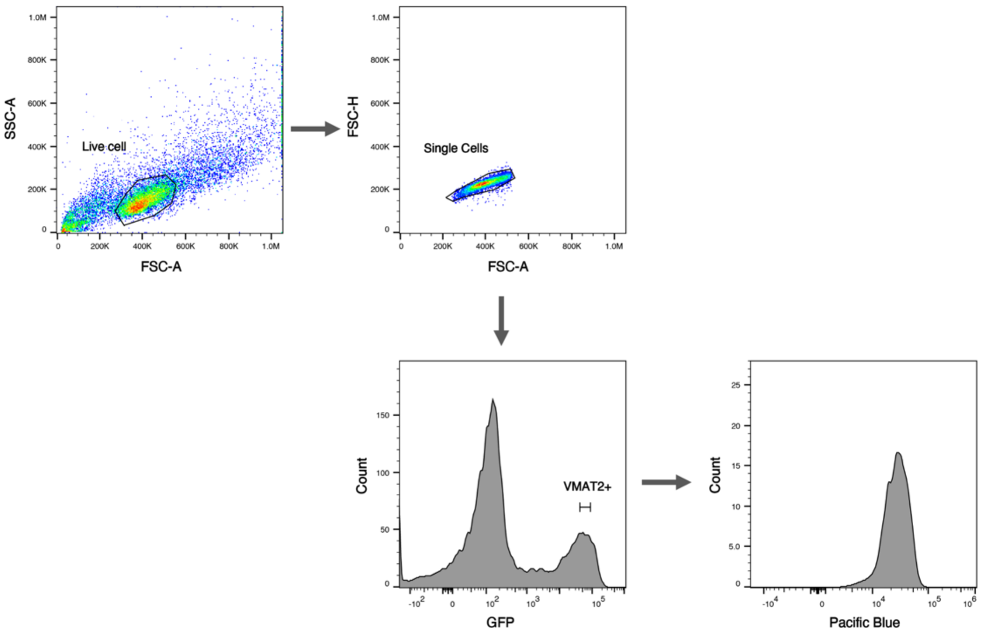


**Fig. S12. Representative flow cytometry gating strategy for the FFN246 uptake assay.** The cells were first gated by FSC and SSC to filter alive cells and single cells. The cells were then gated by GFP fluorescence to select the cells expressing VMAT2 or its mutants. We only selected events which located in a small range of the GFP fluorescence signal to make sure the protein expression level keeping approximately same in the different variants of VMAT2. The selected $GFP^+$ subpopulation was then analyzed for FFN246 uptake level by violet laser channel 405/450.

| | VMAT2A | VMAT2S | VMAT2T | VMAT2$_{YC}$R | VMAT2R | VMAT2$_{YC}$S |
|---|---|---|---|---|---|---|
| **Data collection and processing** | | | | | | |
| Magnification | 130,000 | 130,000 | 130,000 | 105,000 | 130,000 | 130,000 |
| Voltage (kV) | 300 | 300 | 300 | 300 | 300 | 300 |
| Electron exposure ($e^-$/Å$^2$) | 50 | 50 | 50 | 60 | 50 | 50 |
| Defocus range (μm) | -1.2 to -2.0 | -1.2 to -2.0 | -1.2 to -2.0 | -1.2 to -2.0 | -1.2 to -2.0 | -1.2 to -2.0 |
| Pixel size (Å) | 0.932 | 0.932 | 0.932 | 0.832 | 0.932 | 0.932 |
| Symmetry | C1 | C1 | C1 | C1 | C1 | C1 |
| Initial particle (no.) | 21,827,845 | 10,943,366 | 21,933,726 | 9,162,145 | 8,579, 107 | 5,939,244 |
| Final particle (no.) | 220,731 | 124,189 | 335,823 | 132,817 | 166,931 | 175,647 |
| Map resolution (Å) | 3.60 | 3.57 | 3.37 | 3.74 | 4.75 | 4.10 |
| **Refinement** | | | | | | |
| Initial model used (PDB code) | AlphaFold2 model | 8WLJ | 8WLJ | 8WLJ | | |
| Map sharpening *B* factor (Å$^2$) | 202.5 | 199.2 | 200.3 | 217.6 | 385.8 | 274.8 |
| Model composition | | | | | | |
| non-hydrogen atoms | 4,035 | 2,804 | 2,873 | 1,815 | | |
| Protein residues | 530 | 382 | 383 | 389 | | |
| Ligands | | 1 | 1 | 1 | | |
| *B* factor (Å$^2$) | | | | | | |
| Protein | 117.55 | 109.41 | 95.79 | 101.62 | | |
| Ligand | | 100.51 | 91.78 | 82.44 | | |
| R.m.s. deviations | | | | | | |
| Bond lengths (Å) | 0.004 | 0.004 | 0.004 | 0.005 | | |
| Bond angles (°) | 0.762 | 0.789 | 0.856 | 0.940 | | |
| Validation | | | | | | |
| MolProbity score | 1.50 | 1.74 | 1.62 | 1.63 | | |
| Clash score | 7.96 | 9.84 | 9.42 | 8.90 | | |
| Poor rotamers (%) | 0.00 | 0.00 | 0.00 | 0.00 | | |
| Ramachandran plot | | | | | | |
| Favored (%) | 97.71 | 96.56 | 97.36 | 97.14 | | |
| Allowed (%) | 2.29 | 3.44 | 2.64 | 2.86 | | |
| Outliers (%) | 0.00 | 0.00 | 0.00 | 0.00 | | |
| Deposited model | 8WLJ | 8WLM | 8WLK | 8WLL | | |
| Deposited map | EMD-37621 | EMD-37624 | EMD-37622 | EMD-37623 | EMD-37623 | EMD-37623 |

**Table S1. Cryo-EM data collection and refinement statistics.**

| Systems | Protein conformation | Substrate | No. of POPC | No. of Water | No. of Na+ | No. of Cl- | Total atoms | Replicas | Simulation Time (ns) |
|---|---|---|---|---|---|---|---|---|---|
| VMAT2-5HT | Outward | 5HT | 120 | 10527 | 26 | 33 | 53686 | 3 | 500 |
| VMAT2-TBZ | Occluded | TBZ | 120 | 10545 | 27 | 31 | 53750 | 3 | 500 |
| VMAT2-RES | Inward | RES | 118 | 10428 | 27 | 32 | 53296 | 3 | 500 |
| VMAT2-5HT (D399-Protonated) | Outward | 5HT | 120 | 10223 | 26 | 34 | 52777 | 3 | 500 |
| VMAT2-5HT (E312-D399-Protonated) | Outward | 5HT | 120 | 10212 | 26 | 35 | 52638 | 3 | 500 |

**Table S2. Parameters for MD simulation system setup.**
